# Post-stroke reorganization of gait-related functional muscle networks

**DOI:** 10.64898/2026.09.26.752563

**Authors:** Sora Ohnishi, Yusaku Takamura, David O’Reilly, Naomichi Mizuta, Takumi Kai, Mitsuru Konita, Shu Morioka

**Affiliations:** Neurorehabilitation Research Center, Kio University, Nara, Japan; Department of Neurorehabilitation Laboratory, Graduate School of Health Sciences, Kio University, Nara, Japan; School of Biomedical Sciences, University of Leeds, Leeds, United Kingdom; NeuroLandscape, Suwalska 8/78, Warszawa, 03-252, Poland; Department of Rehabilitation, Faculty of Health Sciences, Nihon Fukushi University, Aichi, Japan; Department of Rehabilitation Medicine, Hatsudai Rehabilitation Hospital, Tokyo, Japan; Department of Rehabilitation Medicine, Funabashi Municipal Rehabilitation Hospital, Chiba, Japan

## Abstract

Post-stroke gait impairment is associated with altered muscle coordination. Understanding how functional muscle coordination contributes to joint control is essential for clarifying the mechanisms of gait impairment after stroke. Functional muscle network approaches quantify how muscle interactions contribute to joint kinematics. We investigated how gait-related functional muscle networks are reorganized after stroke and how this reorganization relates to motor impairment and gait dysfunction. Thirty-five participants with subacute stroke and 25 healthy controls underwent gait analysis using surface electromyography and lower-limb kinematics. We quantified functionally similar (redundant information), complementary (synergistic information), and independent (unique information) muscle interactions underlying joint kinematics. Participants with stroke exhibited greater redundant information and lower independent information than healthy controls. Stroke-specific functional muscle network modules emerged through merging modules observed in healthy controls. Stroke-specific module activation was also associated with motor impairment and gait performance. Overall, these findings identify a shift toward more redundant muscle coordination and stroke-specific reorganization of functional network modules, providing new insights into post-stroke gait dysfunction.

## INTRODUCTION

Human gait depends on precise coordination across multiple muscles and joints to support stable, adaptable locomotion [1,2]. Such coordination enables the neuromuscular system to control the many degrees of freedom of the musculoskeletal system while producing efficient movement. After stroke, motor paralysis, abnormal muscle tone, and impaired selective motor control disrupt neuromuscular coordination and contribute to gait dysfunction and reduced walking ability [2–5]. Muscle synergies have been proposed as a mechanism by which the central nervous system coordinates the activation of multiple muscles during movement. Accordingly, muscle synergy analysis characterizes muscle coordination as low-dimensional modules and has been widely used to study coordination during walking [6,7]. Previous studies have reported fewer muscle synergies and altered coordination patterns after stroke, suggesting reorganization of modular coordination [8–11].

The conventional muscle-synergy framework is based on the motor redundancy problem, in which the numerous muscular degrees of freedom are controlled through common neural inputs [12,13]. Accordingly, muscle synergies are extracted from the temporal and spatial correlations in muscle activity, and primarily identify correlated activation patterns reflecting shared neural input across multiple muscles [6,7]. Although this framework has provided important insights into motor coordination, how different types of muscle interactions contribute to joint control during post-stroke gait remains incompletely understood. Recent work suggests that motor control depends not only on common neural inputs but also on complementary interactions among elements with distinct functional roles and contributions [14–17]. For example, anatomically adjacent muscles with similar biomechanical functions receive partially shared neural inputs while exhibiting distinct patterns of coordination with other muscles, suggesting differentiated functional contributions to motor control [18,19]. Furthermore, complementary interactions among multiple elements can generate information that emerges only from their joint activity and cannot be explained by any single element. Such interactions may support flexible motor coordination by integrating distinct functional information across the motor system [20,21]. However, the distinct functional contributions of these muscle interactions to joint control during post-stroke gait remain unclear.

Recently, the Network Information Framework (NIF) has emerged as a novel approach for examining functional muscle coordination [22,23]. Building on conventional muscle synergy analysis, NIF combines information-theoretic and network-based approaches to characterize distinct types of functional muscle interactions according to their contributions to movement and joint kinematics. Within this framework, partial information decomposition (PID) decomposes the information provided by multiple elements about a target variable into redundant information shared across elements, independent information contributed by individual elements, and synergistic information that emerges only when multiple elements are considered together [24,25]. These components represent, respectively, overlapping, muscle-specific, and complementary contributions to joint kinematics [15,26]. By characterizing muscle activity in terms of functional contributions to joint kinematics rather than patterns of coactivation or covariation, NIF may provide a novel perspective on the reorganization of muscle coordination after stroke. Furthermore, because gait coordination is inherently spatiotemporal, examining both spatial and temporal functional muscle networks may further clarify post-stroke muscle coordination.

This reorganization of muscle coordination may be associated with impaired selective motor control after stroke [27–29]. Impaired selective motor control may increase reliance on common neural drive and reduce differentiation among individual muscle contributions to joint control during walking. Moreover, stroke-related corticospinal tract dysfunction has been associated with increased reliance on compensatory descending pathways, such as the reticulospinal tract, which may result in more widespread common neural drive across muscle groups [30–32]. Accordingly, post-stroke gait may be characterized by increased redundant information, reduced independent muscle contributions, and less differentiated joint-specific control. Whether these networks are systematically reorganized after stroke and whether such reorganization is associated with motor impairment severity and gait dysfunction remain unknown. Therefore, we hypothesized that functional muscle networks underlying gait are reorganized after stroke and that these alterations are associated with motor impairment severity and gait dysfunction.

We investigated how functional muscle networks underlying gait are reorganized after stroke using the NIF. We decomposed information shared among muscle activations about gait-related joint kinematics into redundant, synergistic, and independent information components and characterized network reorganization through changes in module fractionation and merging. We then examined whether these alterations were associated with motor impairment severity and gait dysfunction. We found that post-stroke gait showed increased redundant information, reduced independent information, and stroke-specific module reorganization through the merging of healthy modules and reduced joint-specific differentiation. These network alterations were associated with motor impairment and gait performance. By characterizing post-stroke gait from the perspective of functional muscle networks, this study provides new insights into the functional neuromuscular organization underlying post-stroke gait.

## RESULTS

### Participant characteristics

Table 1 summarizes the participant characteristics and clinical assessment results. Thirty-five participants with subacute stroke and 25 age-matched healthy controls participated in this study. The mean age was 57.3 ± 11.0 years in the stroke group and 54.3 ± 10.8 years in the healthy control group. Mean age did not differ significantly between groups (*p* = 0.310). Participants with stroke had mean Fugl–Meyer Assessment lower-extremity synergy scores (FMS) and Fugl–Meyer Assessment (FMA) of the lower extremity scores of 15.8 ± 4.2 and 24.7 ± 5.5, respectively, whereas the mean Trunk Impairment Scale (TIS) and Short Form Berg Balance Scale (SFBBS) scores were 19.1 ± 3.3 and 22.6 ± 5.4, respectively. Participants with stroke had a mean gait speed of 0.61 ± 0.25 m/s, stance and swing time ratios of 0.91 ± 0.10 and 1.27 ± 0.28, respectively, and a Functional Ambulation Category (FAC) score of 3.13 ± 0.93. Based on FMS scores, 19 participants were classified as having mild-to-moderate impairment (FMS ≥ 17) and 16 as having severe impairment (FMS ≤ 16).

**Table 1.** Clinical evaluations between groups.

|  | Healthy<br>(n = 25) | Stroke<br>(n = 35) | p-value |
| --- | --- | --- | --- |
| Age (years) | $54.3 \pm 10.8$ | $57.3 \pm 11.0$ | 0.310 |
| Sex (n): male / female | 15 / 10 | 21 / 14 | 1.000 |
| Affected side (n):<br>right / left |  | 18 / 17 |  |
| Stroke type (n):<br>hemorrhage / infarction |  | 21 / 14 |  |
| Time since stroke (days) | | $64.8 \pm 20.2$ | |
| Single-point cane use (n):<br>Yes / No |  | 20 / 15 |  |
| FMS (max 22) | | $15.8 \pm 4.2$ | |
| FMA (max 34) | | $24.7 \pm 5.5$ | |
| MAS (max 4) | | $0.79 \pm 0.7$ | |
| TIS (max 23) | | $19.1 \pm 3.3$ | |
| SFBBS (max 28) | | $22.6 \pm 5.4$ | |
| FAC (max 5) | | $3.13 \pm 0.93$ | |
| Gait speed (m/s) | | $0.61 \pm 0.25$ | |
| Stance time ratio | | $0.91 \pm 0.10$ | |
| Swing time ratio | | $1.27 \pm 0.28$ | |

Values are presented as mean ± SD or number of participants. FMS, Fugl–Meyer Assessment lower-extremity synergy scores; FMA, Fugl–Meyer Assessment; MAS, Modified Ashworth Scale; TIS, trunk impairment scale; SFBBS, short-form Berg balance scale; FAC, functional ambulation category.

### Characteristics of muscle activity and joint kinematics in participants with stroke

Figure 1 presents the time-series profiles of muscle activity and joint kinematics and group comparisons of full width at half maximum (FWHM) and range of motion (ROM) (Fig. 1C). Linear mixed-effects models showed significant group effects on FWHM for the vastus medialis (VM), rectus femoris (RF), lateral hamstrings (LH), medial hamstrings (MH), gastrocnemius (Gas), and soleus (Sol) (all *p* < 0.05). Post hoc analyses showed that FWHM for the VM was greater in the non-paretic limb than in healthy controls (*p* = 0.025). FWHM for the RF was greater in the non-paretic limb than in both healthy controls and the paretic limb (both *p* < 0.05). FWHM values for the LH and MH were greater in both the paretic and non-paretic limbs than in healthy controls (both *p* < 0.001). FWHM for the Gas was greater in the paretic limb than in both healthy controls and the non-paretic limb (both *p* < 0.05), whereas FWHM for the Sol was lower in the non-paretic limb than in both healthy controls and the paretic limb (both *p* < 0.05). Group effects on ROM were significant for the ankle, knee, hip, and limb angles (all *p* < 0.001). These patterns are consistent with previous studies reporting prolonged muscle activation and reduced lower-limb joint ROM during post-stroke gait [33].

**Figure 1.**
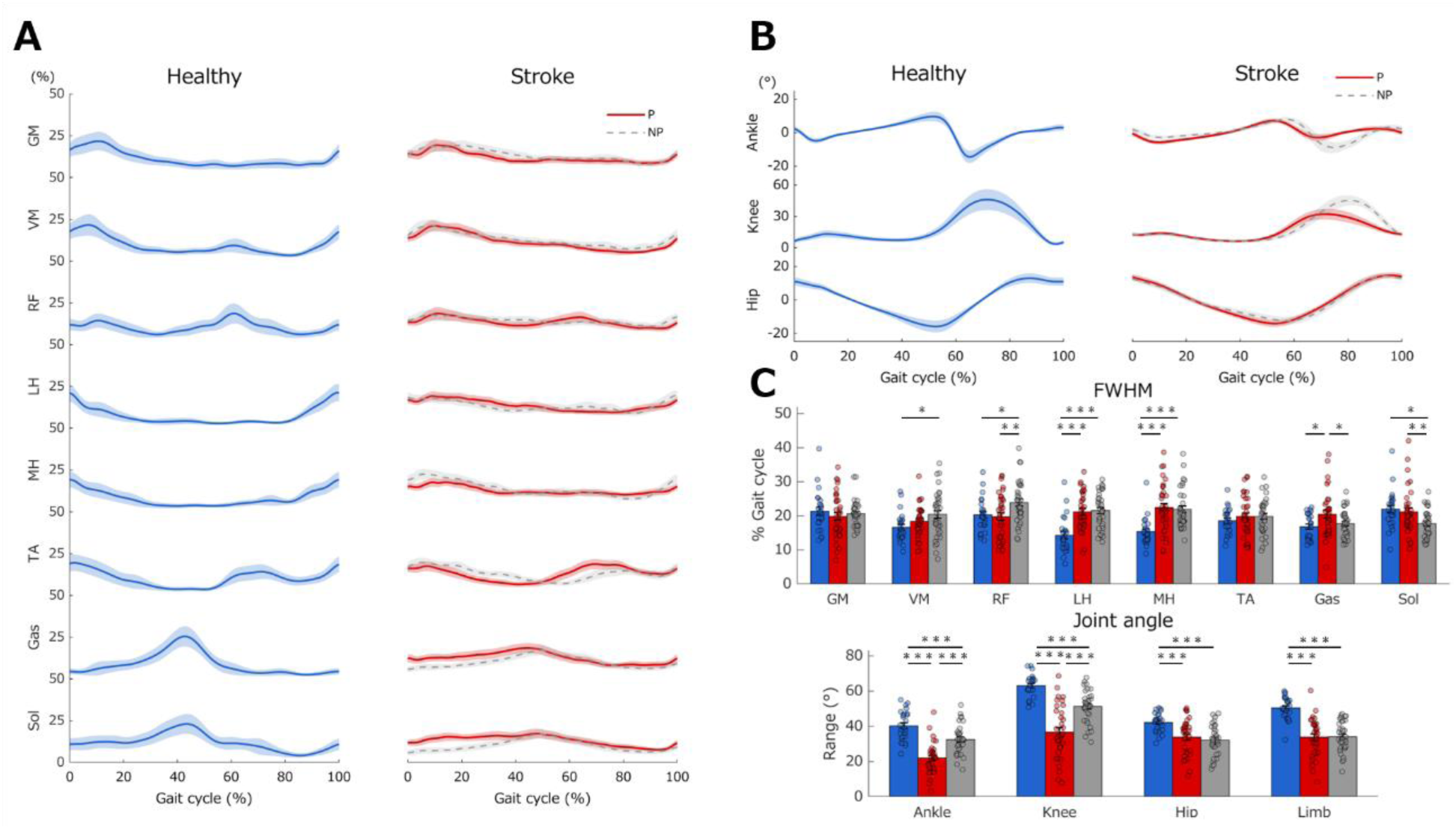
Muscle activity and joint kinematics during gait Time-series profiles of muscle activity (A) and joint kinematics (B), and group comparisons of FWHM and ROM values during gait (C). Blue denotes healthy controls, red denotes the paretic limb of participants with stroke, and gray indicates the non-paretic limb. Waveforms are presented as mean ± 95% confidence intervals. \**p* < 0.05, \*\**p* < 0.01, \*\*\**p* < 0.001. FWHM, full width at half maximum; ROM, range of motion; GM, gluteus medius; VM, vastus medialis; RF, rectus femoris; LH, lateral hamstrings; MH, medial hamstrings; TA, tibialis anterior; Gas, gastrocnemius; Sol, soleus.

### Post-stroke functional muscle network structure shifts toward redundancy

Figure 2 shows group differences in the redundant, synergistic, and independent components, and associations with clinical parameters. A mixed-effects aligned rank transform analysis of variance (ART-ANOVA) showed a significant group × component interaction (F[4, 223.10] = 5.12; *p* = 0.0006; ηp^2^ = 0.084). Post hoc analyses showed that both the paretic and non-paretic limbs of participants with stroke had significantly greater redundant information (*p* = 0.036 and *p* = 0.042, respectively) and lower independent information (*p* = 0.024 and *p* = 0.004, respectively) than the corresponding limbs of healthy controls, whereas no significant group differences were observed for synergistic information (*p* = 0.894 and *p* = 0.545, respectively) (Fig. 2A). To further evaluate evidence for the absence of a group difference in synergistic information, supplementary Bayesian independent-samples t-tests were performed. Bayesian analysis provided moderate support for the null hypothesis when comparing healthy controls and the paretic limb (BF₀₁ = 3.76), whereas evidence for the comparison with the non-paretic limb was inconclusive (BF₀₁ = 0.99). Together, these results indicate that post-stroke gait was characterized by increased redundant and reduced independent information, whereas synergistic information appeared to be relatively preserved in the paretic limb.

**Figure 2.**
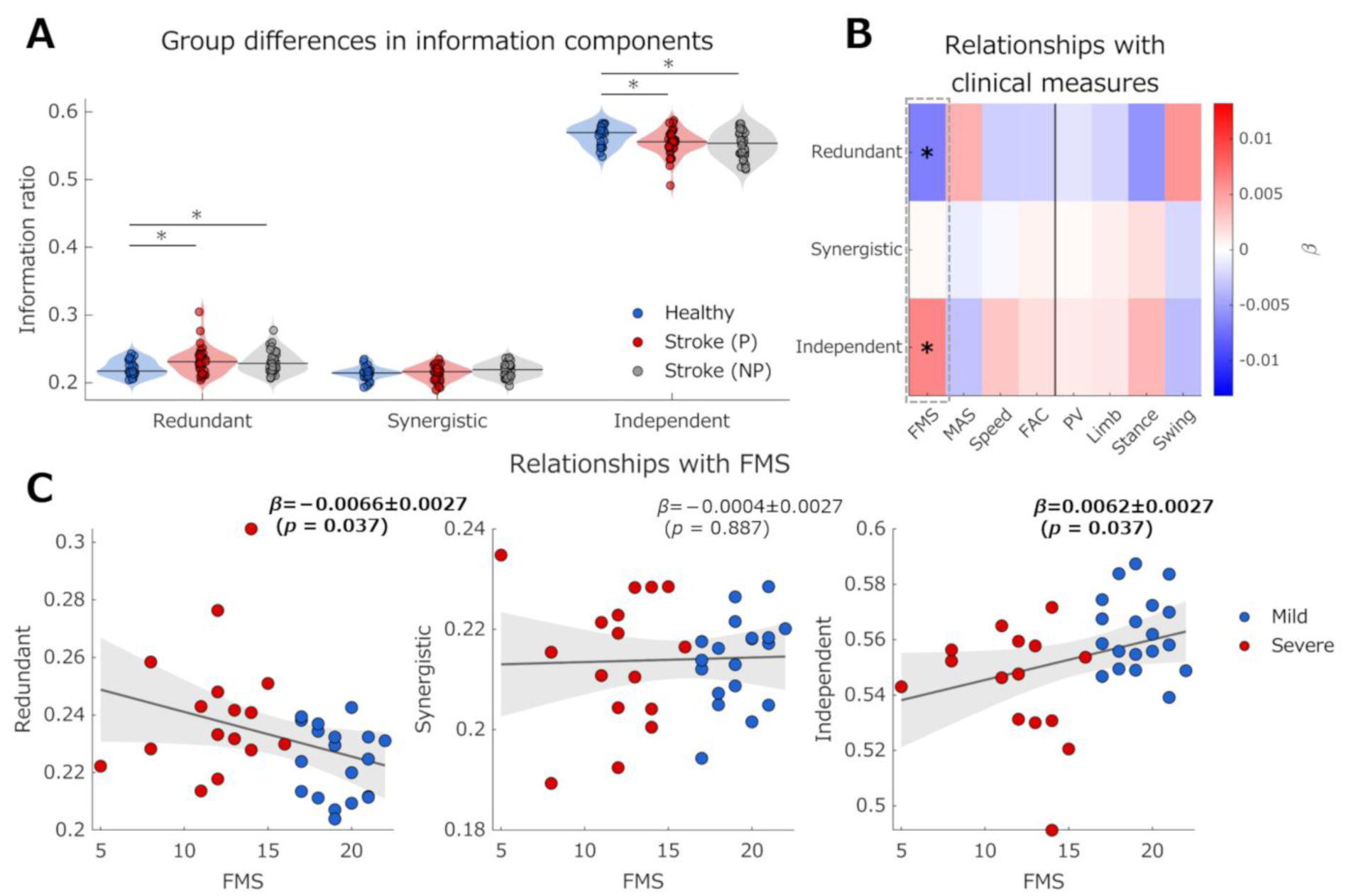
Information components of the functional muscle network and their relationships with clinical parameters (A) Group differences in redundant, synergistic, and independent information components during gait. Blue indicates healthy controls, red indicates the paretic limb of participants with stroke, and gray indicates the non-paretic limb. The central horizontal line within each violin plot indicates the median. (B) Heatmap of regression coefficients (β) from mixed-effects models examining associations between information components and clinical and gait-related measures. Associations with gait-related measures (PV, Limb, Stance, and Swing) were adjusted for FMS. (C) Associations between information components and FMS. Blue indicates the mild subgroup and red indicates the severe subgroup. Solid lines and shaded areas represent the fitted regression lines and 95% confidence intervals, respectively. In panel A, *Holm-adjusted *p* < 0.05. In panels B and C, *FDR-adjusted *p* < 0.05. FDR, false discovery rate; FMS, Fugl–Meyer Assessment lower-extremity synergy scores; MAS, Modified Ashworth Scale; FAC, Functional Ambulation Category; PV, planar covariation; Limb, limb angle range of motion; Stance, stance time ratio; Swing, swing time ratio.

Mixed-effects models showed that higher FMS scores were associated with lower redundant information (β = −0.0066 ± 0.0027; *p* = 0.037) and greater independent information (β = 0.0062 ± 0.0027; *p* = 0.037), whereas no significant association was observed for synergistic information (β = −0.0004 ± 0.0027; *p* = 0.887) (Fig. 2B–C). Thus, greater motor impairment was associated with a shift toward greater redundant and lower independent information.

To examine whether clinical and gait-related associations differed according to motor impairment severity, we characterized severity-dependent differences in information structure using linear mixed-effects models. A significant Information Component × Severity interaction (F[2, 99] = 12.69; p = 0.0001; ηp² = 0.201) reflected greater redundant and lower independent information in the severe than in the mild subgroup (*p* = 0.002 and *p* = 0.002, respectively). Accordingly, subsequent severity-dependent analyses were restricted to the redundant and independent components. A significant Metric × Severity interaction was observed for the association between redundant information and FAC (F[1, 31] = 4.93; *p* = 0.041; ηp² = 0.137) (Supplementary Table 1). Simple-slope analysis showed a positive association in the severe subgroup (β = 0.013 ± 0.0060; *p* = 0.041), but no corresponding association in the mild subgroup (β = −0.0028 ± 0.0040; *p* = 0.486), suggesting that greater redundant information was associated with higher FAC scores only among participants with severe motor impairment.

Figure 3 presents the functional muscle network structure together with results from the threshold-free cluster enhancement (TFCE)-based network-based statistic (NBS). Distinct functional muscle networks were identified for the redundant, synergistic, and independent information components. TFCE-based NBS revealed significantly stronger redundant connectivity between the VM and LH in participants with stroke (*p* = 0.045), whereas significantly stronger independent connectivity between the gluteus medius (GM) and RF was observed in healthy controls (*p* = 0.048) (Fig. 3B). In the non-paretic limb (Supplementary Fig. 1), healthy controls also showed significantly stronger independent connectivity between the GM and Sol muscles (*p* = 0.045). Thus, group differences in functional muscle network topology were spatially localized, involving specific redundant and independent muscle interactions.

**Figure 3.**
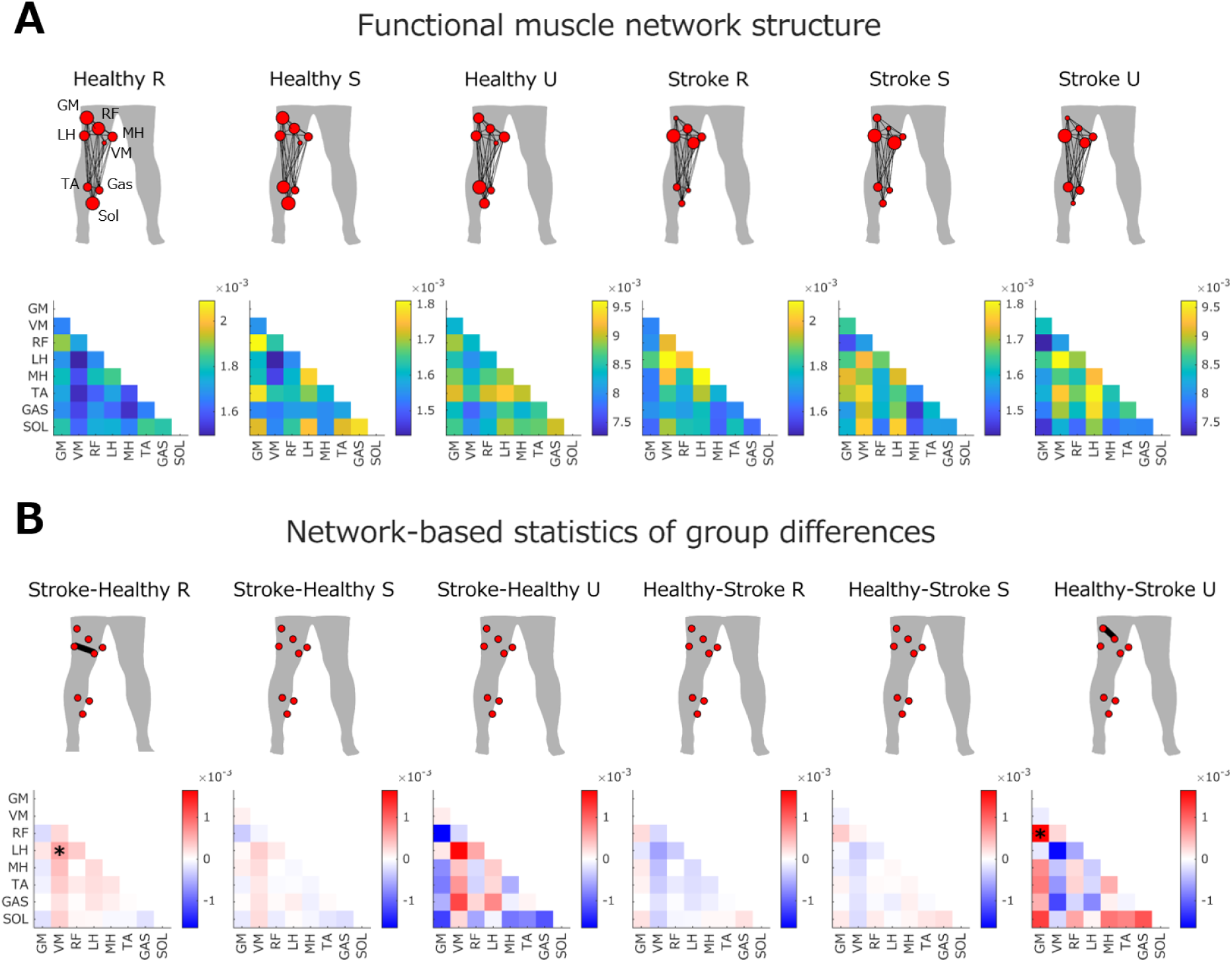
Functional muscle network structure in the paretic limb and threshold-free cluster enhancement-based network statistics (A) Functional muscle network structure of the paretic limb for redundant (R), synergistic (S), and independent (U) information components during gait. Node size indicates the relative strength of each muscle contribution, and line thickness indicates connectivity strength between muscles. Connectivity matrices represent pairwise information strength between muscles. (B) TFCE-based NBS comparison of functional muscle network structure between healthy controls and participants with stroke. Significant connections identified by TFCE-based NBS are indicated by thick black lines in the network graphs. Connectivity matrices represent the difference in pairwise information strength between groups for each information component. Red indicates greater connectivity strength in the first group of the comparison, whereas blue indicates greater connectivity strength in the second group. \**p* < 0.05. GM, gluteus medius; VM, vastus medialis; RF, rectus femoris; LH, lateral hamstrings; MH, medial hamstrings; TA, tibialis anterior; Gas, gastrocnemius; Sol, soleus.

### Post-stroke functional muscle network reorganization is characterized by merging of healthy modules

Functional muscle network modules identified using non-negative matrix tri-factorization (NM3F) differed across groups (Supplementary Figs. 2–4). At group level, five redundant and synergistic modules and four independent modules were identified in healthy controls (Supplementary Fig. 2), compared with four redundant modules and three synergistic and independent modules in the paretic limb (Supplementary Fig. 3), and three redundant, five synergistic, and three independent modules in the non-paretic limb (Supplementary Fig. 4). Participant-level analyses showed significant group differences in module number for all three information components (redundant: F[2, 92] = 3.70; *p* = 0.028; ηp² = 0.075; synergistic: F[2, 92] = 5.35; *p* = 0.011; ηp² = 0.118; independent: F[2, 92] = 7.87; p = 0.0013; ηp² = 0.152). Post hoc comparisons showed fewer modules in the paretic limb than in healthy controls for all three information components (all *p* < 0.05), whereas synergistic and independent modules were also fewer in the paretic than in the non-paretic limb (both p < 0.01).

The post-stroke network included preserved modules resembling healthy organization as well as distinct stroke-specific modules. Stroke-specific modules with low similarity to those of healthy controls were identified across all information components in the paretic limb (Fig. 4A). Redundant space module 1 consisted primarily of the GM, thigh muscles, and tibialis anterior (TA). This module was associated with time module 1 during early stance for all joints, whereas the knee and hip joints were associated with time module 2 during the early swing phase. Synergistic space module 1 consisted primarily of the hamstrings and TA and was associated with time module 2 during stance across all joints. Independent space module 3 consisted primarily of the Gas and Sol. This module was associated with time module 1 during the late stance phase at the ankle joint, whereas the knee and hip joints were associated with time module 2 during the stance and late swing phases. In the non-paretic limb, redundant space module 2 was specific to participants with stroke (Supplementary Fig. 5). This module consisted primarily of the hamstrings and TA. At the ankle, time modules 1 and 3 were associated with the stance and late swing phases, respectively. The knee joint was associated with time module 1 during the stance phase, whereas the hip joint was associated with time module 3 during the late swing phase.

**Figure 4.**
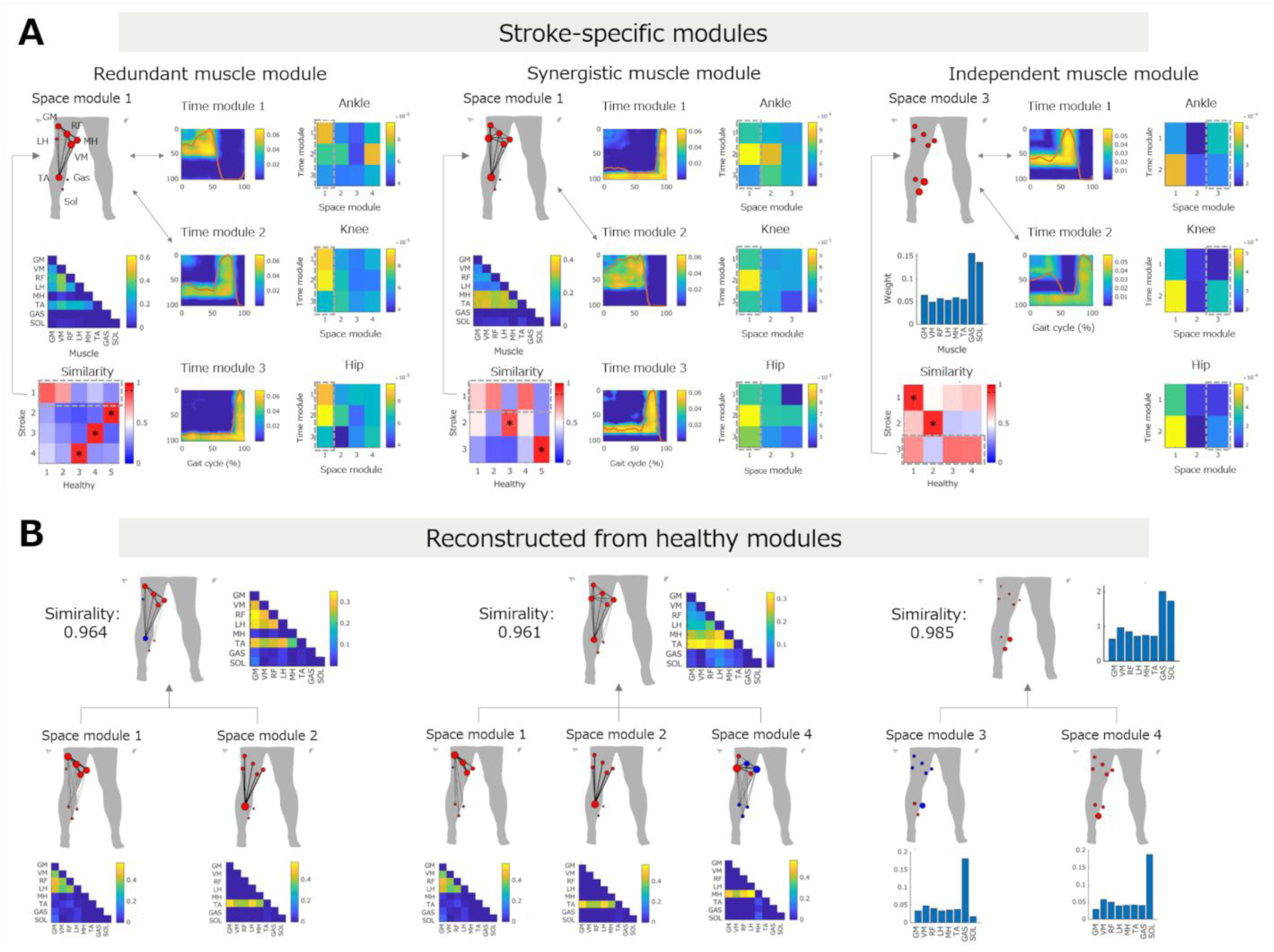
Representative stroke-specific functional muscle network modules in the paretic limb and their reconstruction from healthy modules (A) Representative stroke-specific modules with low similarity to healthy controls are presented for the redundant, synergistic, and independent information components. Similarity matrices represent cosine similarity between modules extracted from healthy controls and participants with stroke, and dashed boxes indicate stroke-specific modules. Connectivity matrices of space modules represent pairwise information strength among muscles. Time module matrices represent the temporal strength of information connectivity across the gait cycle. Heatmaps on the right show connectivity strength between spatial and temporal modules for each joint (ankle, knee, and hip), and arrows mark the spatial–temporal module pairs with the strongest connectivity. Similarity thresholds were 0.89 for redundant modules, 0.87 for synergistic modules, and 0.88 for independent modules. \**p* < 0.05. (B) Stroke-specific spatial modules were reconstructed from representative healthy spatial modules using non-negative least squares. The reconstructed modules showed high similarity to the corresponding stroke-specific modules, indicating that stroke-specific modules can be explained by merging representative healthy modules. Similarity values indicate the cosine similarity between the reconstructed and stroke-specific spatial modules. GM, gluteus medius; VM, vastus medialis; RF, rectus femoris; LH, lateral hamstrings; MH, medial hamstrings; TA, tibialis anterior; Gas, gastrocnemius; Sol, soleus.

Non-negative least squares (NNLS) reconstruction analysis showed that each stroke-specific spatial module could be reconstructed with high similarity from combinations of multiple representative healthy modules (Fig. 4B), supporting module merging as a contributor to the emergence of stroke-specific functional muscle network modules. Separate analyses in the mild and severe subgroups showed subgroup-specific patterns of module merging (Supplementary Figs. 6–8). Redundant module merging was observed in both subgroups, whereas synergistic module merging was additionally observed in the severe subgroup, suggesting that the pattern of network reorganization may vary with motor impairment severity. To assess the functional relevance of module reorganization, we used mixed-effects models, which showed that a greater number of merged modules was associated with lower planar covariation (PV), an index of intersegmental coordination during gait [34], specifically for redundant information (β = −0.554 ± 0.125; *p* = 0.0008), whereas no significant associations were observed for fractionation (Fig. 5). This result suggests that greater redundant module merging is linked to reduced intersegmental coordination during gait.

**Figure 5.**
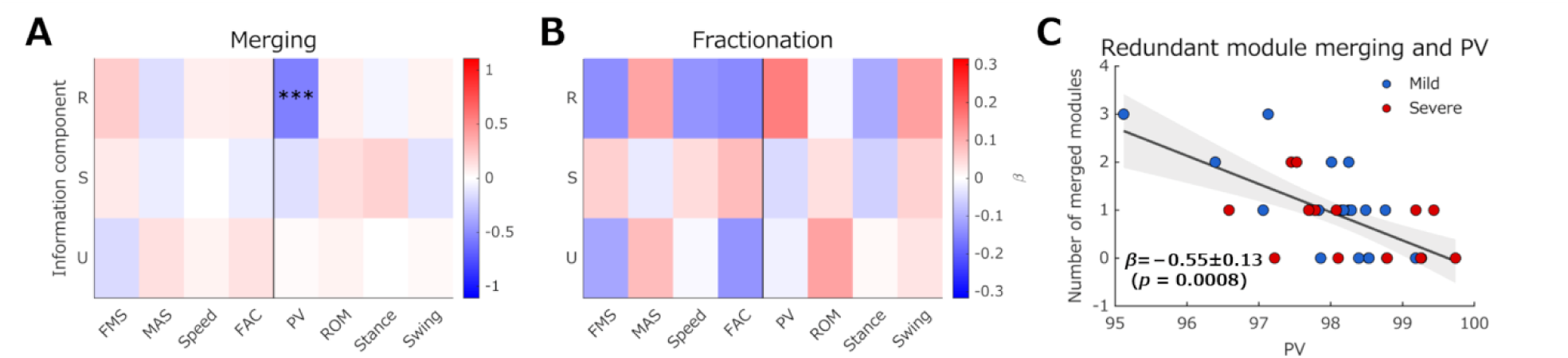
Associations between module reorganization and clinical parameters (A) Heatmap showing regression coefficients (β) from mixed-effects models examining associations between the number of merged modules and clinical and gait-related measures for each information component. Associations with gait-related measures (PV, ROM, Stance, and Swing) were adjusted for FMS. (B) Heatmap showing regression coefficients (β) from mixed-effects models examining associations between the number of fractionated modules and clinical and gait-related measures for each information component. Associations with gait-related measures were adjusted for FMS. (C) Relationship between the number of redundant merged modules and PV. Blue indicates the mild subgroup and red indicates the severe subgroup. The solid line and shaded area represent the fitted regression line and 95% confidence interval, respectively. Regression coefficients (β) are presented as estimates ± standard errors. In panel A, ***FDR-adjusted *p* < 0.001. FDR, false discovery rate; FMS, Fugl–Meyer Assessment lower-extremity synergy scores; MAS, Modified Ashworth Scale; FAC, Functional Ambulation Category; PV, planar covariation; ROM, limb angle range of motion; Stance, stance time ratio; Swing, swing time ratio.

### Post-stroke functional muscle network modules exhibit reduced joint specificity

Comparisons of joint contributions revealed reduced joint specificity in participants with stroke (Fig. 6). In healthy controls, several modules exhibited joint-specific contributions, including a greater hip joint contribution in redundant space module 4, a greater ankle joint contribution in redundant space module 5, a greater knee joint contribution in synergistic space module 3, a greater ankle joint contribution in synergistic space module 5, and a greater knee joint contribution in independent space module 3 (all *p* < 0.05). By contrast, participants with stroke showed fewer joint-specific differences across modules, with only the synergistic space module 2 showing a greater contribution from the ankle joint than from the knee joint (*p* < 0.05). In the non-paretic limb (Supplementary Fig. 9), redundant space module 1 showed a greater contribution from the ankle joint than from the hip joint, whereas synergistic space module 3 had a greater contribution from the hip joint than from the knee joint (both *p* < 0.05).

**Figure 6.**
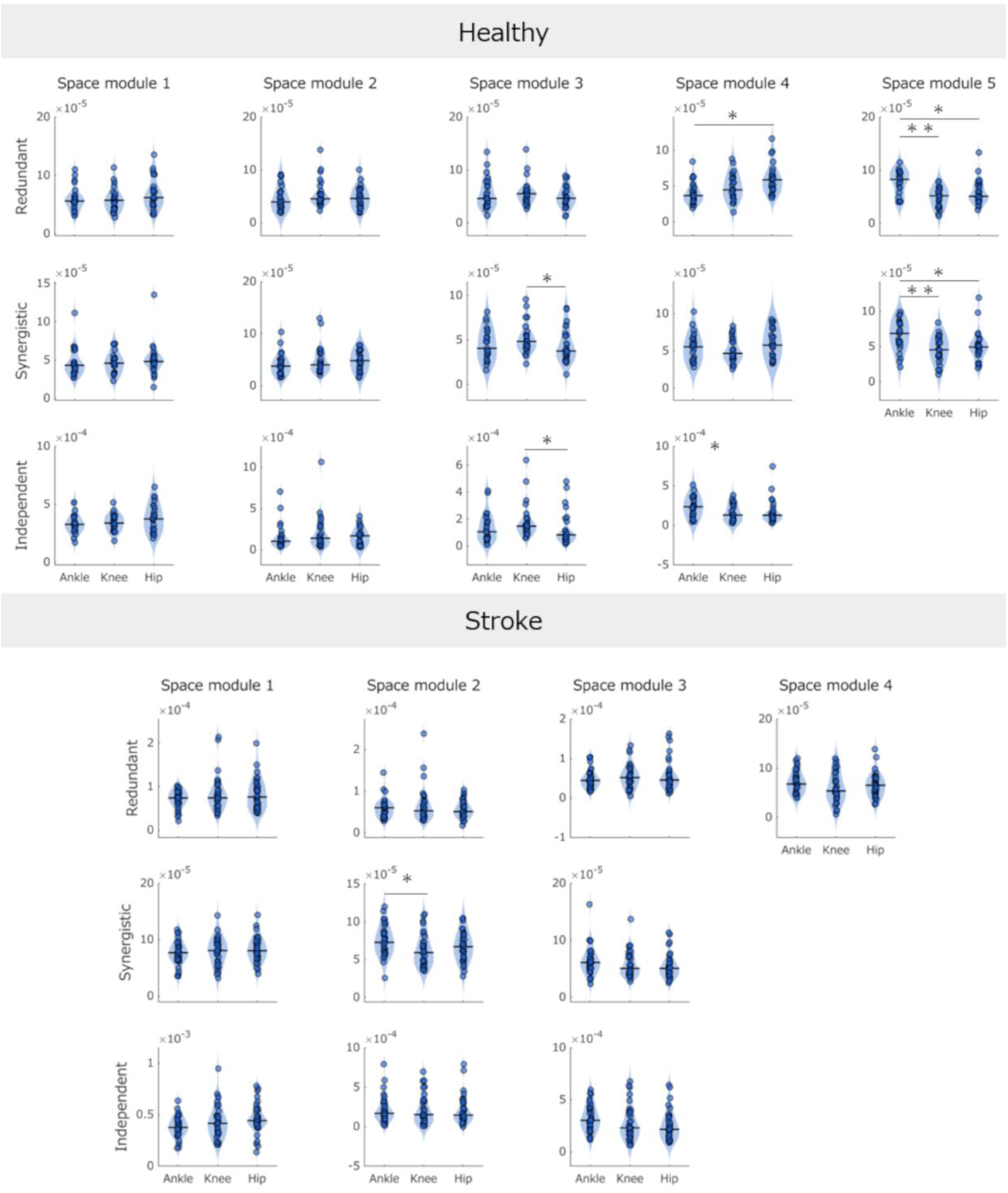
Joint-specific contributions of functional muscle network modules in healthy controls and participants with stroke Comparisons of joint contributions across redundant, synergistic, and independent space modules in healthy controls and participants with stroke. Violin plots show the distribution of joint-specific contributions for the ankle, knee, and hip joints within each space module. Horizontal black lines denote median values. *Holm-adjusted *p* < 0.05, **Holm-adjusted *p* < 0.01.

Severity-specific analyses using mixed-effects ART-ANOVA (Supplementary Fig. 10) showed significant Joint × Severity interactions for synergistic space module 2 (F[2, 66] = 3.20; *p* = 0.042; ηp^2^ = 0.055), synergistic space module 3 (F[2, 99] = 3.71; *p* = 0.035; ηp^2^ = 0.070), and independent space module 3 (F[2, 99] = 3.01; *p* = 0.045; ηp^2^ = 0.050). Post hoc analyses demonstrated significant joint-specific differences only in the mild subgroup (all *p* < 0.05), whereas no such differences were observed in the severe subgroup, consistent with partial preservation of joint specificity in individuals with milder motor impairment.

### Functional muscle network modules are associated with motor impairment and gait performance

Mixed-effects models showed that several functional muscle network modules, including the stroke-specific modules identified in Fig. 4, were associated with motor impairment and gait-related measures (Fig. 7). Higher activation of redundant module 1 was associated with lower FMS scores (β = [−1.13 ± 0.49] × 10⁻⁵; *p* = 0.044) and slower gait speed (β = [−1.17 ± 0.48] × 10⁻⁵; *p* = 0.035). Higher activation of synergistic module 1 was also associated with lower FMS scores (β = [−0.85 ± 0.34] × 10⁻⁵; *p* = 0.042), whereas its activation was positively associated with PV (β = [1.05 ± 0.34] × 10⁻⁵; *p* = 0.009). Together with the reconstruction results in Fig. 4B, these results indicate that stroke-specific modules characterized by the merging of healthy network structures are functionally relevant to motor impairment and gait organization.

**Figure 7.**
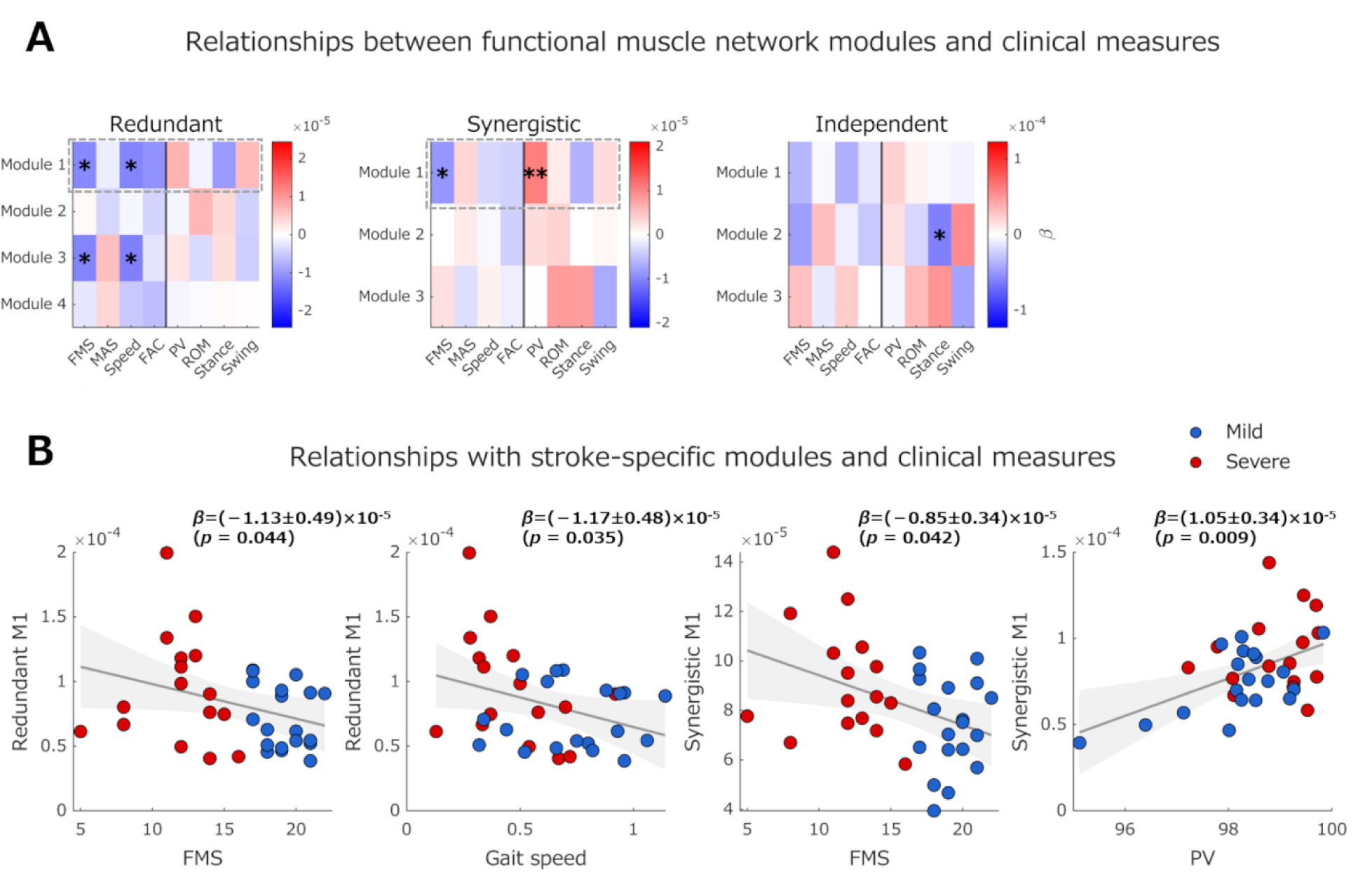
Relationships between functional muscle network modules and clinical parameters (A) Heatmaps of regression coefficients (β) from mixed-effects models examining associations between functional muscle network module strength and clinical and gait-related measures. Associations with gait-related measures (PV, Limb, Stance, and Swing) were adjusted for FMS. (B) Scatter plots of significant associations between functional muscle network modules and selected clinical and gait-related measures. Blue indicates the mild subgroup and red indicates the severe subgroup. Solid lines and shaded areas represent the fitted regression lines and 95% confidence intervals, respectively. Regression coefficients (β) are presented as estimates ± standard errors. In panel A, *FDR-adjusted *p* < 0.05 and **FDR-adjusted *p* < 0.01. FDR, false discovery rate; FMS, Fugl–Meyer Assessment lower-extremity synergy scores; MAS, Modified Ashworth Scale; FAC, Functional Ambulation Category; PV, planar covariation; Limb, limb angle range of motion; Stance, stance time ratio; Swing, swing time ratio.

Severity-dependent differences in functional muscle network modules were examined using mixed-effects Module × Severity models. Significant Module × Severity interactions were identified for redundant (F[3, 132] = 3.37; *p* = 0.020; ηp² = 0.071), synergistic (F[2, 99] = 4.53; *p* = 0.013; ηp² = 0.083), and independent information (F[2, 99] = 3.83; *p* = 0.025; ηp² = 0.071). Post hoc analyses showed greater activation in the severe than in the mild subgroup for redundant modules 1 and 3 (*p* = 0.047 and *p* = 0.014, respectively) and synergistic module 1 (*p* = 0.038) (Supplementary Table 2). Subsequent analyses of these severity-sensitive modules identified Metric × Severity interactions for gait speed and stance ratio in redundant module 1 and for gait speed in redundant module 3 (all *p* < 0.05) (Supplementary Table 3). Follow-up analyses indicated that, in the severe subgroup, redundant module 1 activation was negatively associated with gait speed (β = [−2.27 ± 0.48] × 10⁻⁵; *p* = 0.049) and stance ratio (β = [−2.20 ± 0.48] × 10⁻⁵; *p* = 0.012), whereas redundant module 3 activation was negatively associated with gait speed (β = [−2.37 ± 0.48] × 10⁻⁵; *p* = 0.041); the corresponding associations were not significant in the mild subgroup. These results indicate that the relationship between redundant functional network modules and gait performance depends on motor impairment severity, with significant associations observed only in the severe subgroup.

## DISCUSSION

We assessed motor function, gait kinematics, and muscle activity in healthy controls and participants with subacute stroke and used the NIF to characterize post-stroke reorganization of functional muscle networks during gait. We found that post-stroke gait was characterized by increased redundant and reduced independent information, both of which were associated with motor impairment severity (Fig. 2). Participants with stroke also exhibited fewer functional muscle network modules, stroke-specific coordination patterns that could be reconstructed by merging several healthy modules and were associated with motor impairment and gait performance (Figs. 4 and 7), and reduced joint-specific contributions across modules (Fig. 6). Collectively, these findings indicate that post-stroke gait is characterized by reorganization of functional muscle networks.

Participants with stroke had greater redundant information but lower independent information than healthy controls, and both components were associated with motor impairment severity (Fig. 2). Specifically, redundant information was negatively associated with the FMS score, whereas independent information was positively associated with the FMS score. These results suggest that post-stroke gait control shifts toward a coordination strategy characterized by greater redundancy and reduced independence. Redundant information reflects overlapping information about joint kinematics across multiple muscles. Although overlapping information may enhance the robustness of motor control [20,21,35], excessive redundancy may reduce coordination flexibility. In contrast, independent information reflects the differentiated contributions of individual muscles to joint control. Taken together, increased redundancy and reduced independent information may reflect diminished flexibility and selectivity of motor control after stroke. One possible mechanism for this shift toward greater redundancy and reduced independence is increased reliance on common descending neural drive through compensatory recruitment of alternative descending pathways, such as the reticulospinal pathway, following corticospinal tract damage [30–32]. Such compensation may promote more stable but less differentiated muscle coordination during gait. Notably, although greater redundant information was associated with greater motor impairment overall, its positive association with functional ambulation in the severe subgroup suggests that this coordination strategy may also help preserve walking ability when selective motor control is substantially impaired (Supplementary Table 1). This interpretation is consistent with previous evidence suggesting that alternative descending pathways contribute to motor function after severe stroke [36,37].

This interpretation is consistent with findings from O’Reilly and colleagues, who reported that recovery of upper-limb motor function was accompanied by reduced redundant information and increased synergistic information, suggesting that redundancy may represent an early compensatory coordination strategy [38]. By contrast, overall synergistic information did not differ between groups in the present study, and Bayesian analysis provided moderate evidence supporting the absence of a difference between healthy controls and the paretic limb (BF₀₁ = 3.76). Several factors may explain this discrepancy. The present study examined steady-state walking in a cross-sectional comparison, whereas the previous study investigated upper-limb motor recovery longitudinally. Preserved synergistic information during steady-state walking may reflect interactions between supraspinal and spinal locomotor circuits, with the relative contribution of spinal circuitry being greater during rhythmic locomotion. Because steady-state walking depends more strongly on spinal locomotor networks than many voluntary upper-limb movements [1,39], synergistic information may remain relatively preserved during this task despite corticospinal damage. Locomotor behaviors requiring greater supraspinal involvement, however, such as gait initiation, obstacle negotiation, or turning, may place different demands on complementary muscle interactions [40]. Future work should determine whether synergistic information becomes disrupted during these more adaptable locomotor tasks.

All three information components showed fewer functional muscle network modules and distinct stroke-specific coordination patterns (Fig. 4). Reconstruction analyses further showed that representative stroke-specific modules could be reconstructed by merging several healthy modules, suggesting that post-stroke functional muscle network reorganization may involve the integration of existing coordination modules through module merging. Similar reconstruction patterns emerged after stratification by motor severity, indicating that module merging is a common feature of functional muscle network reorganization across different levels of impairment (Supplementary Figs. 6–8). These findings are consistent with prior studies showing that post-stroke motor coordination is characterized not only by changes in the number of coordination modules but also by module merging, fractionation, and altered muscle weighting [8–11,41,42]. However, in contrast to previous reports, the extent of module reorganization was not consistently associated with conventional clinical measures of motor impairment in our cohort [42,43]. This discrepancy may reflect the subacute stage of recovery in our participants, during which neural reorganization and functional recovery are still evolving. Instead, greater redundant module merging was associated with reduced PV (Fig. 5), a fundamental feature of normal intersegmental coordination during walking. Previous studies have shown that simplified neuromuscular organization is associated with poorer gait kinematics after stroke [44]. These results suggest that greater merging of redundant functional muscle networks is associated with impaired intersegmental coordination during gait.

A similar pattern was observed for synergistic information. Despite the preserved overall magnitude of synergistic information (Fig. 2), synergistic information was represented by fewer modules with stroke-specific organization, suggesting that complementary muscle interactions also undergo modular reorganization. Thus, preservation of the overall amount of synergistic information does not necessarily imply preservation of its modular organization. Examination of stroke-specific modules revealed a common pattern of broad coupling between proximal and distal muscles across multiple joints and gait phases. Unlike the more localized coordination patterns observed in healthy controls, these widespread interactions may reflect reduced selectivity of muscle control and a shift toward more globally coordinated motor strategies after stroke [27–29,45]. This widespread coordination may also reflect compensatory recruitment of proximal musculature and reduced independence of distal muscle control, as reported following stroke [28,45,46].

Consistent with this interpretation, joint-specific contributions were lower across functional muscle network modules (Fig. 6). Whereas healthy controls exhibited joint-specific functional specialization across several modules, participants with stroke showed fewer joint-specific differences. Severity-dependent analyses also showed that joint-specific differences were retained in the mild subgroup but were not evident in the severe subgroup (Supplementary Fig. 10), suggesting that the loss of functional differentiation across joints is more pronounced with greater motor impairment. This pattern is consistent with previous evidence of abnormal inter-joint coupling after stroke and impaired modulation of inter-joint kinematic synergies during gait in individuals with poorer motor recovery [28,47]. Reduced joint specificity may therefore reflect diminished selective control of muscle contributions to individual joint movements, consistent with the broader coupling across muscles and joints observed in the stroke-specific modules. Together, these results suggest that post-stroke functional muscle network reorganization involves not only module merging and the emergence of stroke-specific coordination patterns, but also a loss of joint-specific functional specialization.

Several representative stroke-specific functional muscle network modules were associated with motor impairment severity, gait speed, and stance time ratio (Fig. 7). Consistent with previous studies reporting associations between altered muscle coordination and motor impairment after stroke, our findings further support the relationship between muscle coordination patterns and clinical function [8–11]. In particular, associations with FMS indicate that alterations in functional muscle network organization are closely related to motor impairment severity, whereas associations with gait speed and stance time ratio suggest that these network alterations are linked to functional gait performance. Severity-specific analyses further showed that the relationships between individual functional muscle network modules and clinical outcomes differed according to impairment severity (Supplementary Tables 2 and 3), with most associations observed only in the severe subgroup. Previous muscle synergy studies have similarly suggested that the functional relevance of post-stroke coordination patterns may vary across patient subgroups [43]. Consistent with this notion, our results suggest that the functional significance of specific muscle coordination patterns varies across levels of post-stroke motor impairment. Taken together, these findings indicate that stroke-specific reorganization of functional muscle network modules reflects clinically meaningful changes in motor control underlying post-stroke gait dysfunction.

### Limitations of the study

This study has several limitations. First, its cross-sectional design precluded assessment of longitudinal changes during motor recovery. Second, only muscle activity and joint kinematics were analyzed, and direct neurophysiological measures of descending motor pathways were not obtained. Accordingly, the proposed mechanisms involving altered descending neural drive, including increased reliance on the reticulospinal pathway, remain speculative. Third, the findings were obtained from participants with subacute stroke, and generalizability to other stages of stroke recovery remains uncertain. Finally, walking speed was not recorded in healthy controls, and the use of assistive devices in some participants may have influenced muscle coordination patterns. Future studies integrating neurophysiological measurements with NIF-based analyses could further clarify the neural mechanisms underlying post-stroke gait control.

## CONCLUSIONS

Our findings suggest that post-stroke gait is associated with increased redundant information, reduced independent information, and reorganization of functional muscle network modules linked to motor impairment severity and gait performance. These results provide a new perspective on functional muscle network reorganization after stroke. Future studies should determine whether these findings generalize to other locomotor tasks and to clarify the neural mechanisms underlying functional muscle network reorganization after stroke.

## METHODS

### Participants

This cross-sectional study included 35 participants with subacute stroke (within 6 months of stroke onset) and 25 age-matched healthy controls. Participants with stroke were recruited from Hatsudai Rehabilitation Hospital and Funabashi Municipal Rehabilitation Hospital. Exclusion criteria were: inability to complete the 10-m walking task without physical assistance, inability to walk without a lower-limb orthosis, bilateral brain lesions, Mini-Mental State Examination score less than 24 points, a history of orthopedic disease, pain affecting walking, cerebellar lesions or resting tremor, and unilateral spatial neglect. The study was approved by the Research Ethics Committee of Kio University (Approval No. R7-12). All participants provided written informed consent before study participation.

### Experimental environment and protocol

Participants performed four walking trials along a 10-m walkway with an additional 6-m section to accommodate acceleration and deceleration. All assessments were performed under the supervision of a physical therapist to minimize the risk of falls. Participants could use a cane if required during the walking assessments.

Kinematic and electromyographic data were recorded concurrently during walking. Inertial measurement units (IMUs) (Xsens Technologies B.V., Enschede, The Netherlands; sampling rate, 100 Hz) were placed bilaterally on the feet, shanks, thighs, lumbar region, and upper trunk to measure lower-limb and trunk kinematics. Wireless surface electromyography (EMG) (Delsys Trigno, Delsys Inc., Boston, MA, USA; sampling rate, 2000 Hz) was recorded bilaterally from eight lower-limb muscles (16 muscles in total): GM, VM, RF, LH, MH, TA, Gas, and Sol. For healthy controls, only data from the dominant limb were analyzed. Before electrode placement, the skin was shaved (when necessary) and cleaned with alcohol to reduce skin impedance. Before data collection, participants were confirmed to score <4 on the modified Borg scale to minimize the influence of fatigue on gait performance.

### Clinical evaluation

The FMA [48] was used to assess motor and sensory impairments. The FMS was used to classify participants with stroke into two groups according to motor impairment severity: mild-to-moderate (FMS ≥ 17) and severe (FMS ≤ 16) [5]. Spasticity of the ankle plantar flexors was assessed using the MAS [49,50], with a score of 1+ converted to 1.5 for quantitative analysis. Trunk function, balance, and walking ability were evaluated using the TIS [51], SFBBS [52], and FAC [53], respectively.

### Data analysis

Walking speed was measured with a stopwatch over the central 10 m of the walkway. Gait cycles were identified using sagittal-plane angular velocity data obtained from IMUs attached to both shanks. Initial contact was identified from the angular velocity waveform, after which stance and swing phase durations were calculated. To minimize effects of acceleration and deceleration, the first and last three gait cycles of each trial were excluded. Twenty steady-state gait cycles were obtained from each participant for subsequent analyses.

### IMU data processing

Before acquisition, the IMU sensors were calibrated during quiet standing following the manufacturer’s instructions. Joint kinematic data were filtered using a zero-lag fourth-order Butterworth band-pass filter (0.15–10 Hz). IMU data were used to derive the ankle, knee, hip, and lower-limb segment elevation angles during walking. Ankle joint angles were calculated from the relative orientation between the foot and shank sensors, knee joint angles from the thigh and shank sensors, and hip joint angles from the lumbar and thigh sensors. ROM was defined as the difference between the maximum and minimum joint angles within each gait cycle.

PVs of the lower-limb segment elevation angles were assessed using principal component analysis of the thigh, shank, and foot elevation angles during gait [54]. The analysis was performed separately for each gait cycle, and the percentages of variance explained by the first, second, and third principal components (PV1, PV2, and PV3, respectively) were calculated. The sum of PV1 and PV2 served as an index of intersegmental coordination during gait [34]. Values were then averaged across gait cycles for each participant.

### EMG data processing

The EMG signals were band-pass filtered using a zero-lag fourth-order Butterworth filter (20–500 Hz), de-meaned, full-wave rectified, and low-pass filtered at 10 Hz to obtain the linear envelope. EMG amplitudes were normalized to the maximum value across the extracted gait cycles. EMG preprocessing, including electrode placement, followed the recommendations of the Surface Electromyography for the Non-Invasive Assessment of Muscles guidelines. For PID analysis, each gait cycle was time normalized and resampled to 1,000 samples to preserve temporal resolution for time window-based information analysis before segmentation into temporal windows [23]. For conventional non-negative matrix factorization (NMF)-based muscle synergy analysis, gait cycles were resampled to 200 samples.

Peak EMG activity was defined as the maximum of the time-normalized EMG signal within each gait cycle. The FWHM was calculated as the duration for which EMG activity exceeded 50% of its peak value within each gait cycle, representing the temporal width of muscle activation during gait [34,55]. Peak EMG activity and FWHM values were then averaged across gait cycles for each participant.

### NIF analysis

NIF-based analysis of the functional muscle networks

To characterize functional muscle networks underlying gait, we applied NIF [22,23], which uses PID to quantify functional interactions between muscle activity and joint kinematics. Twenty gait cycles of EMG and joint-angle data were time-normalized to 1,000 samples and segmented into 50 temporal windows of 20 samples each. Corresponding temporal windows were pooled across gait cycles before PID analysis [23,56].

PID is an information-theoretic framework that decomposes the joint mutual information between multiple source variables and a target variable into redundant, unique (independent), and synergistic information components [24,25]. Specifically, PID partitions the joint mutual information, I(m_x_, m_y_; τ), provided by two muscle EMG signals (m_x_ and m_y_) about a target variable (τ) into redundant information shared between muscles (R), unique (independent) information from each muscle (U), and synergistic information that emerges only from complementary interactions between muscles (S) (Equation 1).

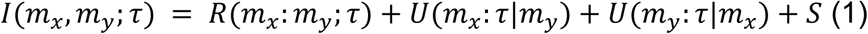

The two unique information terms were averaged to obtain a single independent information value for each muscle pair. In this study, pairwise combinations of muscle EMG signals served as source variables, whereas ankle, knee, and hip joint angles served as target variables. PID was implemented using a Gaussian PID framework based on local information estimates and Gaussian copula mutual information [57] to quantify redundant, synergistic, and independent information linking muscle pairs and joint kinematics.

To enable comparisons across participants and joints, each PID component was normalized by the total mutual information, defined as the sum of redundant, synergistic, and independent information. The normalized PID values were represented as functional adjacency matrices describing the intermuscular functional interactions during walking. For the spatial analysis, PID values were calculated for all pairwise muscle combinations across temporal windows and joint variables, yielding spatial networks that characterized intermuscular functional interactions. For the temporal analysis, PID values were calculated across all pairs of temporal windows for each muscle pair, yielding temporal networks that characterized the time-varying functional interactions throughout the gait cycle. Spatiotemporal functional muscle networks were then constructed by integrating the spatial and temporal information networks, consistent with previous information-theoretic analyses of muscle coordination [23,56].

To retain statistically meaningful functional connections, the network matrices were thresholded using a modified percolation analysis [58,59]. Briefly, network connections were iteratively thresholded based on the percolation threshold estimated from random networks of equivalent size. Thresholding was terminated before fragmentation of the largest connected component (the giant component), which preserved long-range functional connectivity while removing weak or potentially spurious connections [58,59]. Negative values were excluded from further analyses.

To evaluate the relative contribution of each component to the overall information structure of muscle coordination, the proportions of redundant, synergistic, and independent information relative to total mutual information were calculated for each participant. This approach follows prior studies emphasizing the balance among information components as a fundamental characteristic of functional system organization [38].

Extraction of functional muscle network modules

Community detection was applied separately to the spatial and temporal information networks to determine optimal modular organization. For the spatial information networks, overlapping communities were identified using a modified link-based community detection approach [60]. Specifically, complete-linkage hierarchical clustering of network links was used to construct link dendrograms, which were partitioned at the level maximizing partition density. Binary co-membership matrices indicating whether pairs of muscles belonged to the same community were generated for each participant and network layer. These matrices were aggregated across network layers and participants to construct a consensus co-membership graph; link-based community detection was then applied to identify the consensus spatial community structure.

For the temporal information networks, communities were identified using the Louvain algorithm based on modularity maximization [61]. Binary co-membership matrices indicating whether pairs of temporal windows belonged to the same community were generated for each participant and network layer. These matrices were aggregated into a consensus co-membership graph, and the Louvain algorithm was reapplied to identify the consensus temporal community structure. The resulting numbers of spatial and temporal modules were used as input parameters for dimensionality reduction.

To extract functional muscle network modules, NM3F was applied separately to the redundant, synergistic, and independent information matrices [23,56]. The spatiotemporal information matrix was factorized into spatial, temporal, and coupling matrices that described relationships between spatial and temporal modules. Spatial modules represented muscle coordination patterns, whereas temporal modules represented gait phase-dependent information patterns. Coupling matrices quantified associations between spatial and temporal modules for each joint. Joint contribution matrices were used to quantify the relative contributions of each spatial module to ankle, knee, and hip joint control. For analyses of clinical associations, module activation coefficients were averaged across temporal modules and joint variables to obtain a representative activation coefficient for each spatial module. This approach evaluated the overall clinical relevance of spatial muscle coordination patterns rather than phase-or joint-specific effects.

### Subnetwork analysis

To quantify the relative importance of individual muscles within the extracted functional muscle network modules, the total communicability of each node C(i) was computed [62]. Total communicability was defined as the row-wise sum of the matrix exponential of the adjacency matrix (A), and reflected the extent to which a node communicates with all other nodes in the network by accounting for walks of all lengths (Equation 2).

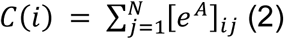

To visualize the most salient functional connections within each module, connections with communicability values below the network mean were omitted from the human-body visualizations. To identify salient subnetworks of functionally related muscles, the link-based community detection approach described above was applied to each extracted module.

### Identification of common and stroke-specific modules

To distinguish common from stroke-specific modules, cosine similarity was calculated between the module weight vectors derived from both conventional NMF and NIF-based analyses [63]. For each module identified in the stroke group, maximum cosine similarity with the corresponding modules in the healthy control group was calculated. The median of these maximum similarity values was defined as the observed similarity index. To estimate the chance-level threshold, the module weight vectors were randomly permuted across muscles, and cosine similarities between the healthy control and stroke modules were recalculated. For each permutation, the maximum similarity between each stroke module and the corresponding healthy modules was calculated, and the median of these maximum similarity values was recorded. This procedure was repeated 10,000 times to generate a null distribution of similarity indices. The 95th percentile of the null distribution defined the significance threshold for module similarity. Modules with maximum cosine similarity exceeding this threshold were classified as common modules, whereas those below the threshold were classified as stroke-specific modules (non-preserved) [64].

These stroke-specific modules were analyzed using an NNLS-based reconstruction procedure adapted from Cheung et al. [42] to characterize module reorganization. Stroke modules were reconstructed from healthy modules to identify module merging, whereas healthy modules were reconstructed from stroke modules to identify module fractionation. Modules reconstructed with at least two contributing modules (NNLS coefficient > 0.20) and a reconstruction cosine similarity of 0.75 or more were classified as merged or fractionated modules. When a stroke module satisfied both criteria, it was assigned to the category with the higher reconstruction cosine similarity. For each participant and information component, the numbers of merged and fractionated modules were calculated as the respective counts of modules classified as merged or fractionated. These counts were used to examine associations between module reorganization and clinical and gait-related parameters.

### Statistics and Reproducibility

Group differences in joint kinematics, muscle activation characteristics, conventional muscle synergy measures, and the number of functional muscle network modules were analyzed using linear mixed-effects models, with group as a fixed effect and participant as a random intercept. Group differences in normalized information components were evaluated using mixed-effects ART-ANOVA, with group and information component as fixed effects and participant as a random intercept. For non-significant inter-group comparisons, Bayesian independent-samples *t*-tests were also performed to quantify evidence for the null hypothesis using the default Jeffreys–Zellner–Siow prior (*r* = 0.707), and Bayes factors (BF₀₁) were reported. Joint-specific contributions of extracted modules were assessed using Friedman tests, whereas severity-dependent differences were evaluated using mixed-effects ART-ANOVA with joint, severity, and their interaction (Joint × Severity) as fixed effects and participant as a random intercept.

Associations with clinical (FMS, MAS, gait speed, and FAC) and gait-related measures (PV, ROM, and stance and swing phase time ratios) were analyzed using linear mixed-effects models. Models included the standardized metric and its interaction with information component or module as fixed effects and participant as a random intercept; module analyses were performed separately for each information component. Component-and module-specific regression coefficients (β) and standard errors were estimated from these models. The same framework was used to assess associations between the numbers of merged or fractionated modules and clinical and gait-related measures. For gait-related measures, FMS was included as an additional covariate.

Severity-dependent analyses were performed in two stages. First, differences by impairment severity were evaluated using linear mixed-effects models including Information Component × Severity or Module × Severity, with participant as a random intercept, followed by post hoc comparisons between the mild and severe subgroups. Second, for components or modules showing severity-dependent differences, linear models including the standardized metric, severity, and their interaction (Metric × Severity) were used to test whether associations differed by severity. Significant interactions were followed by simple-slope analyses within each subgroup.

Multiple testing was controlled using the Benjamini–Hochberg FDR procedure across information components for each metric and across modules within each information component and metric. The same correction was applied across information components for merging and fractionation analyses. The Holm correction was applied to post hoc pairwise comparisons.

To characterize local-network alterations in functional muscle interactions, TFCE-based NBS was applied to compare the functional muscle networks corresponding to the redundant, synergistic, and independent information components between healthy controls and participants with stroke [65,66]. Two-sample t-statistics were calculated for corresponding network edges between groups, and significance was assessed by permutation testing (5,000 permutations) with family-wise error rate correction. TFCE enhanced detection of spatially clustered network differences without requiring an arbitrary cluster-forming threshold, thereby identifying clusters of altered functional muscle interactions between groups.

Statistical significance was set at p < 0.05. All analyses were conducted in MATLAB R2021b (MathWorks, Natick, MA, USA). MATLAB code for the NIF methodology is publicly available from *the* DelisLab/Muscle_PID GitHub repository.

## DATA AVAILABILITY

The datasets generated and/or analyzed in this study are available from the corresponding author upon reasonable request.

### Code availability

Code and example data are available at the following repository: https://github.com/DelisLab/Muscle_PID.

## ACKNOWLEDGEMENTS

We thank the staff of the Kio University Neuro-Rehabilitation Research Centre for their advice and support throughout the study. We also thank the clinical staff of Hatsudai Rehabilitation Hospital and Funabashi Municipal Rehabilitation Hospital for their assistance with participant recruitment and data collection.

## AUTHOR CONTRIBUTIONS

Conceptualization, S.O., Y.T., and S.M.; Methodology, S.O., Y.T., and D.O.; Investigation, S.O., T.K., and M.K.; Formal analysis, S.O. and D.O.; Data curation, S.O.; Visualization, S.O.; Interpretation, S.O., Y.T., D.O., and N.M.; Writing – original draft, S.O.; Writing – review & editing, all authors; Funding acquisition, S.M.; Project administration, S.M.; Supervision, Y.T. and S.M. All authors read and approved the final manuscript.

## ETHICS APPROVAL

This study was approved by the Research Ethics Committee of Kio University (Approval No. R7-12).

## INFORMED CONSENT

Written informed consent was obtained from all participants before study participation.

## SUPPLEMENTARY INFORMATION

Supplemental methods, Supplementary Fig. 1–12, and Supplementary Tables 1–3.

