## SUPPLEMENTARY INFORMATION for "Post-stroke reorganization of gait-related functional muscle networks"

### ***Conventional muscle synergy analysis***

For conventional muscle synergy analysis, time-normalized EMG signals were concatenated across gait cycles without averaging across cycles, as previously described [1]. NMF was applied separately to each participant to extract coordinated muscle activation patterns termed muscle synergies [2]. NMF decomposed the EMG matrix as follows:

$EMG = (W_{m\times n} * C_{n\times t}) + error$ (Supplementary Equation 1)

where W represents the synergy weight matrix and the relative contribution of each muscle to each synergy, whereas C contains the temporal activation coefficients describing the activation of each synergy across the concatenated gait cycles. NMF was implemented with a multiplicative update algorithm. Convergence tolerances for residual error and relative changes in matrix elements were set to $10^{-6}$ and $10^{-4}$, respectively. The algorithm was run 1,000 times, and the solution yielding the lowest root-mean-square residual error was selected [3]. Each synergy weight vector was normalized by its maximum value to enable comparisons across participants and conditions. The optimal number of modules was defined as the minimum number of modules explaining >90% of the total variance accounted for (VAF) in the original EMG signals. The variance accounted for by the first module (VAF1) was also calculated as an index of the extent to which overall muscle activity could be represented by a single dominant synergy [4].

Synergy structures were compared across participants and groups using hierarchical clustering using the synergy weight vectors [5,6]. The optimal cluster number was determined from the average silhouette score, and the cluster number with the highest local silhouette value was selected [7,8].

For each representative module, the highest cosine similarity between the representative module and the synergies extracted from an individual participant defined the participant-specific module similarity score. These scores were used to evaluate associations with clinical and gait-related parameters.

Conventional NMF-based muscle synergy analysis showed significantly greater VAF1 in both the paretic and non-paretic limbs than in healthy controls (both $p < 0.001$), with no group differences in the number or overall structure of muscle synergies (Supplementary Fig. 11).

Associations between NMF-derived modules and clinical and gait-related parameters are presented in Supplementary Fig. 12. Significant associations were identified between module 1 and FAC ($\beta = 0.044 \pm 0.023$, $p = 0.046$), module 2 and stance time ratio ($\beta = 0.046 \pm 0.022$, $p = 0.045$), and module 4 and swing time ratio ($\beta = 0.078 \pm 0.023$, $p = 0.003$). Module similarity was not significantly associated with FMS. No significant Module × Severity interaction was detected ($F[3, 132] = 0.74$, $p = 0.527$, $\eta p^2 = 0.016$), providing no evidence for severity-dependent differences in NMF-derived module associations.

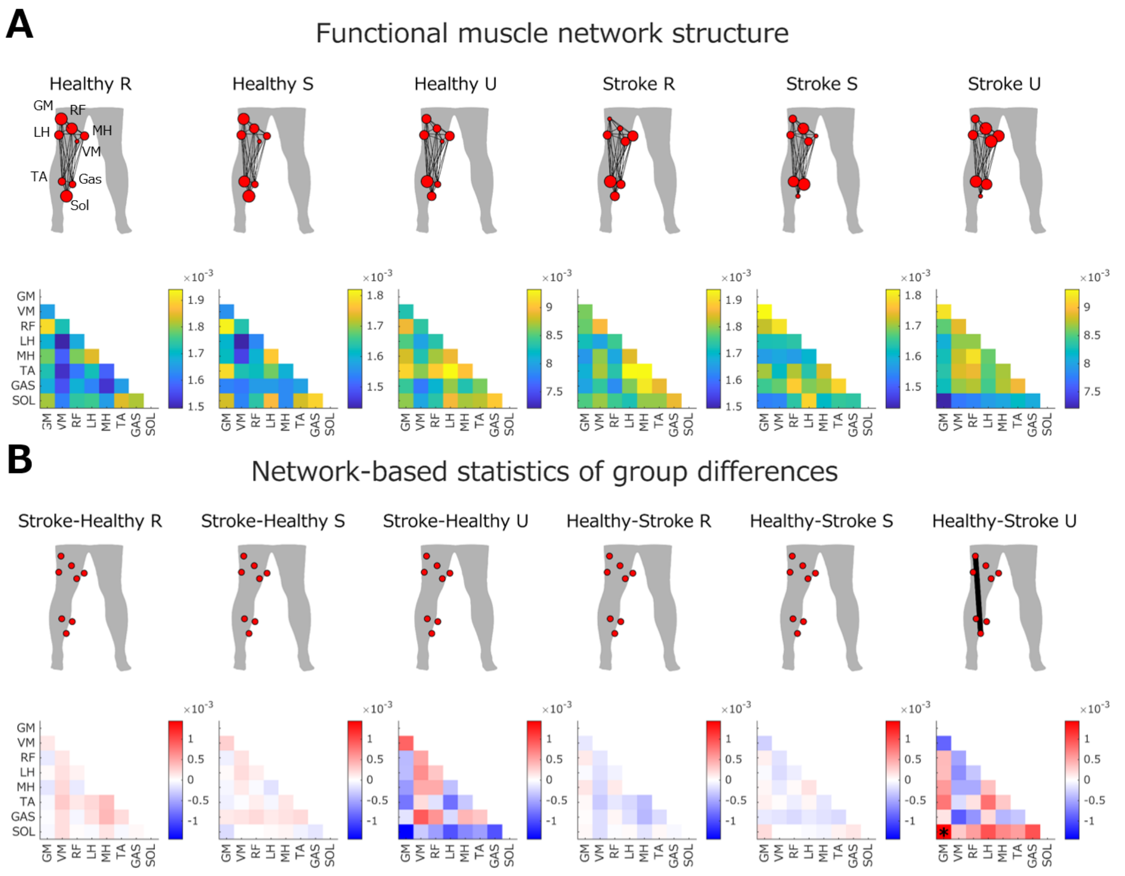


Supplementary Fig. 1. Functional muscle network structure in the non-paretic limb and threshold-free cluster enhancement-based network statistics

(A) Functional muscle network structure of the non-paretic limb for redundant (R), synergistic (S), and independent (U) information components during gait. Node size indicates the relative strength of each muscle contribution, and line thickness indicates connectivity strength between muscles. Connectivity matrices show pairwise information strength between muscles. (B) TFCE-based NBS comparison of functional muscle network structure between healthy controls and participants with stroke. Connectivity matrices show differences in pairwise information strength between groups for each information component. Red indicates greater connectivity strength in the first group of the comparison, whereas blue indicates greater connectivity strength in the second group. *$p < 0.05$. GM, gluteus medius; VM, vastus medialis; RF, rectus femoris; LH, lateral hamstrings; MH, medial hamstrings; TA, tibialis anterior; Gas, gastrocnemius; Sol, soleus.

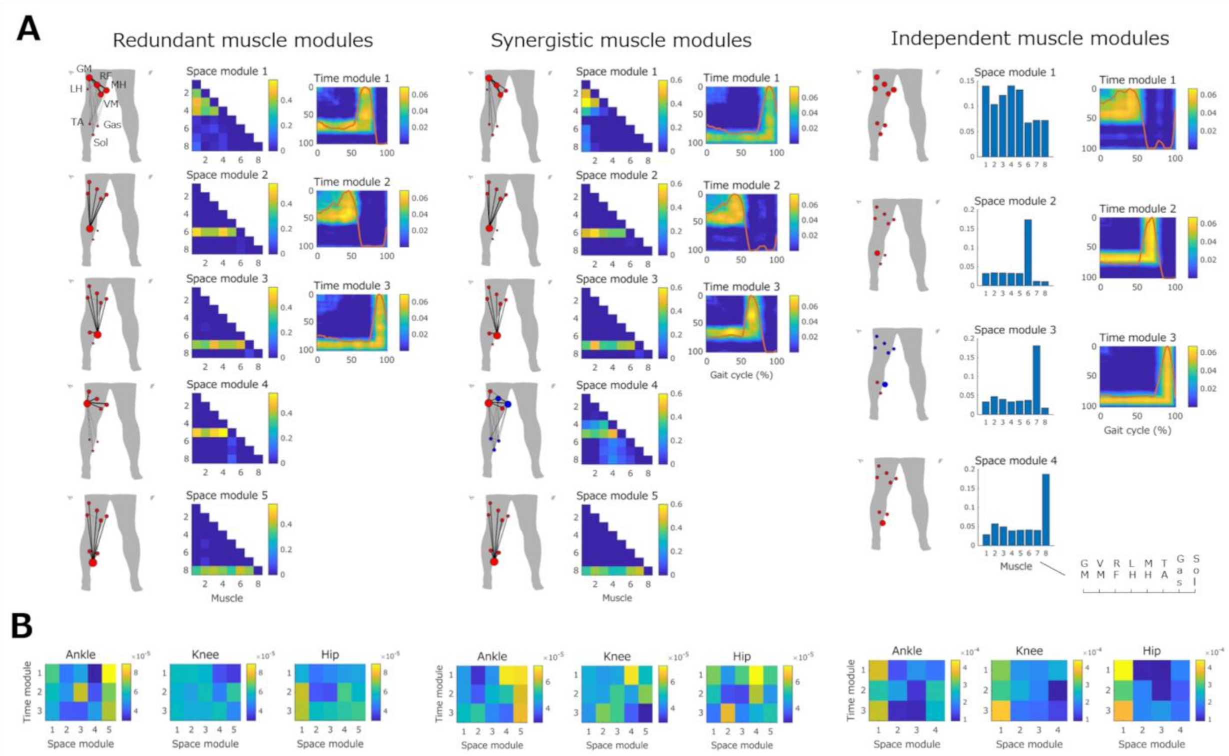


Supplementary Fig. 2. Functional muscle network modules in healthy controls

(A) Functional muscle network modules extracted from healthy controls. Node size indicates the relative strength of each muscle contribution, and line thickness indicates connectivity strength among muscles. Colored node groups denote subnetworks identified using link-based community detection. Space-module connectivity matrices show pairwise information strength among muscles. Time-module matrices show the temporal strength of information connectivity across the gait cycle. Muscle order was identical across all spatial modules: GM, VM, RF, LH, MH, TA, Gas, and Sol. (B) Relationships between space modules and time modules for each joint in healthy controls. Connectivity matrices show coupling strength between space and time modules for each joint. Individual space modules were associated with multiple time modules depending on joint-specific functional contributions. GM, gluteus medius; VM, vastus medialis; RF, rectus femoris; LH, lateral hamstrings; MH, medial hamstrings; TA, tibialis anterior; Gas, gastrocnemius; Sol, soleus.

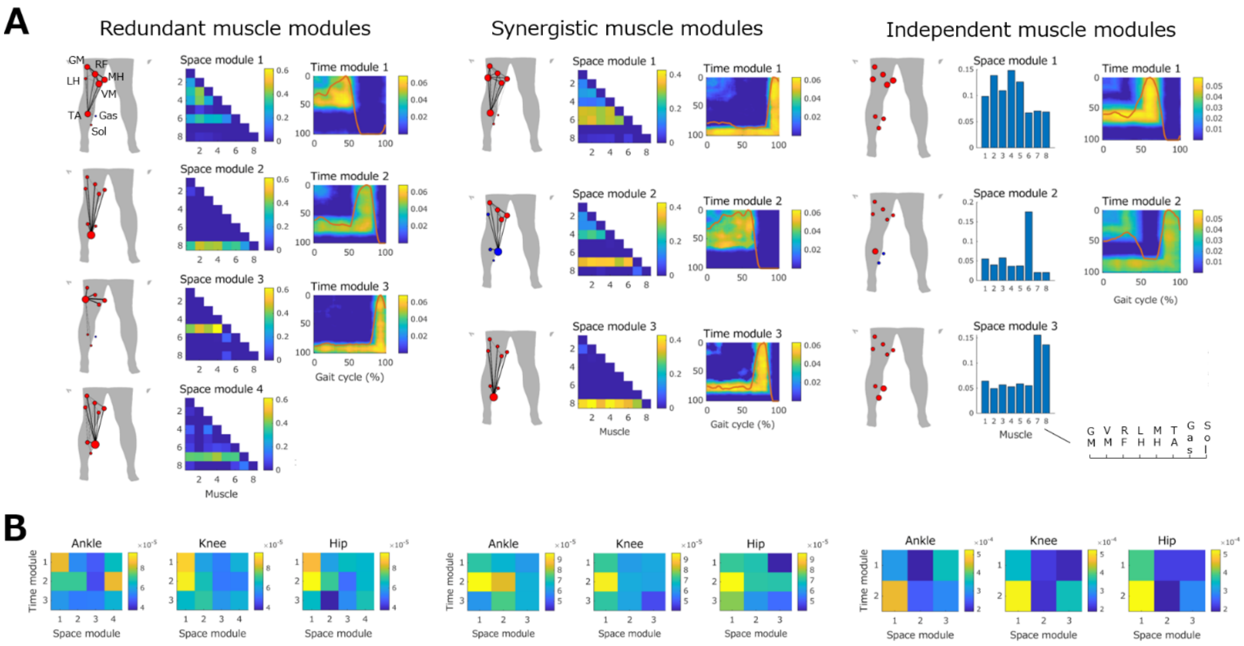


Supplementary Fig. 3. Functional muscle network modules in the paretic limb of participants with stroke

(A) Functional muscle network modules extracted from the paretic limb of participants with stroke. Node size indicates the relative strength of each muscle contribution, and line thickness indicates connectivity strength among muscles. Colored node groups denote subnetworks identified using link-based community detection. Space-module connectivity matrices show pairwise information strength among muscles. Time-module matrices show the temporal strength of information connectivity across the gait cycle. Muscle order was identical across all spatial modules: GM, VM, RF, LH, MH, TA, Gas, and Sol. (B) Relationships between space modules and time modules for each joint in the paretic limb of participants with stroke. Connectivity matrices show coupling strength between space and time modules for each joint. Individual space modules were associated with multiple time modules depending on joint-specific functional contributions. GM, gluteus medius; VM, vastus medialis; RF, rectus femoris; LH, lateral hamstrings; MH, medial hamstrings; TA, tibialis anterior; Gas, gastrocnemius; Sol, soleus.

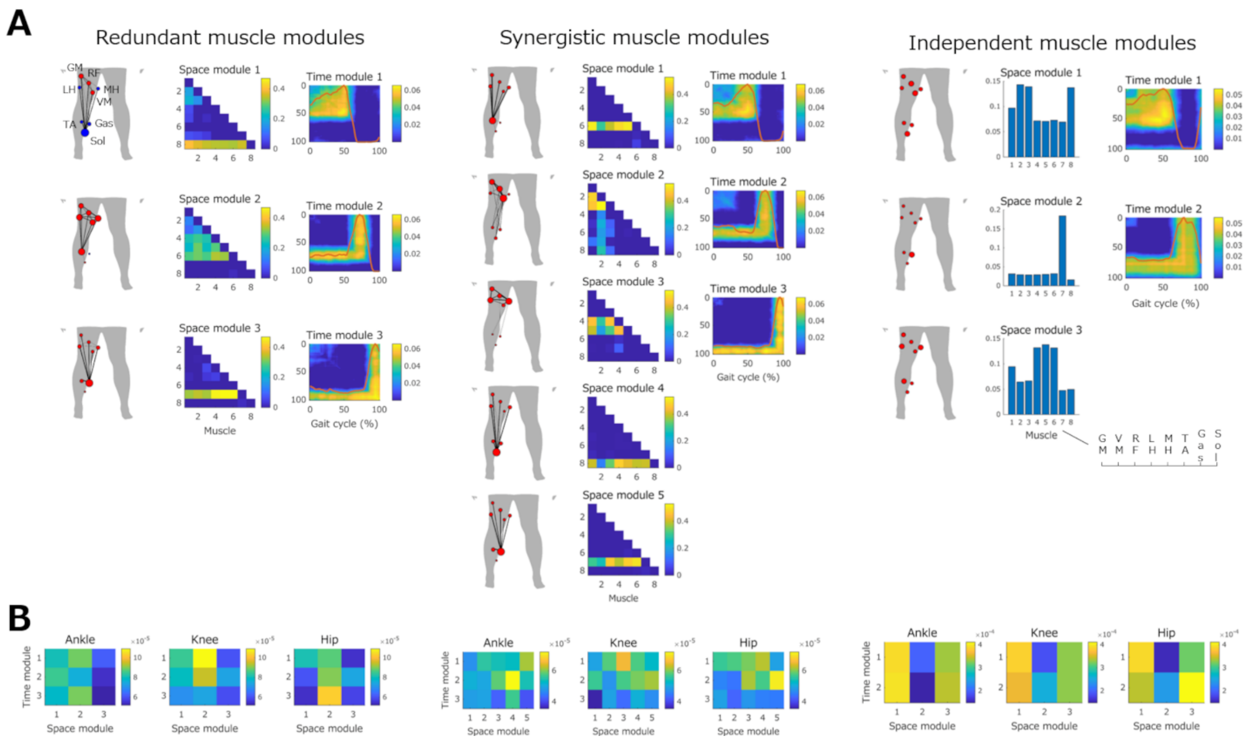


Supplementary Fig. 4. Functional muscle network modules in the non-paretic limb of participants with stroke

(A) Functional muscle network modules extracted from the non-paretic limb of participants with stroke. Node size indicates the relative strength of each muscle contribution, and line thickness indicates connectivity strength among muscles. Colored node groups denote subnetworks identified using link-based community detection. Space-module connectivity matrices show pairwise information strength among muscles. Time-module matrices show the temporal strength of information connectivity across the gait cycle. Muscle order was identical across all spatial modules: GM, VM, RF, LH, MH, TA, Gas, and Sol. (B) Relationships between space modules and time modules for each joint in the non-paretic limb of participants with stroke. Connectivity matrices show coupling strength between space and time modules for each joint. Individual space modules were associated with multiple time modules depending on joint-specific functional contributions. GM, gluteus medius; VM, vastus medialis; RF, rectus femoris; LH, lateral hamstrings; MH, medial hamstrings; TA, tibialis anterior; Gas, gastrocnemius; Sol, soleus.

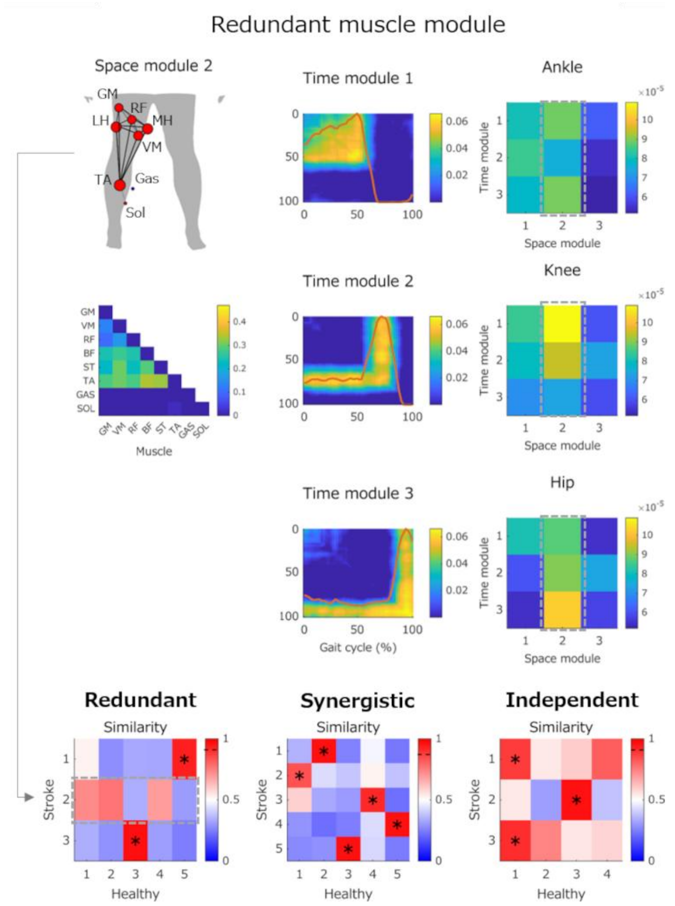


Supplementary Fig. 5. Representative stroke-specific functional muscle network module in the non-paretic limb

Representative redundant functional muscle network module with low similarity to healthy controls in the non-paretic limb of participants with stroke. Similarity matrices represent cosine similarity between modules extracted from healthy controls and the non-paretic limb of participants with stroke. Dashed boxes indicate the stroke-specific module with low similarity to healthy controls selected for further analysis. Among all modules, only redundant space module 2 showed low similarity to healthy controls, whereas the remaining modules were highly similar to those in healthy controls. Space-module connectivity matrices show pairwise information strength among muscles, whereas time-module matrices show the temporal strength of information connectivity across the gait cycle. Arrows between space and time modules indicate strong spatiotemporal associations between modules. The similarity thresholds were 0.89 for redundant modules, 0.87 for synergistic modules, and 0.89 for independent modules. *p <

0.05. GM, gluteus medius; VM, vastus medialis; RF, rectus femoris; LH, lateral hamstrings; MH, medial hamstrings; TA, tibialis anterior; Gas, gastrocnemius; Sol, soleus.

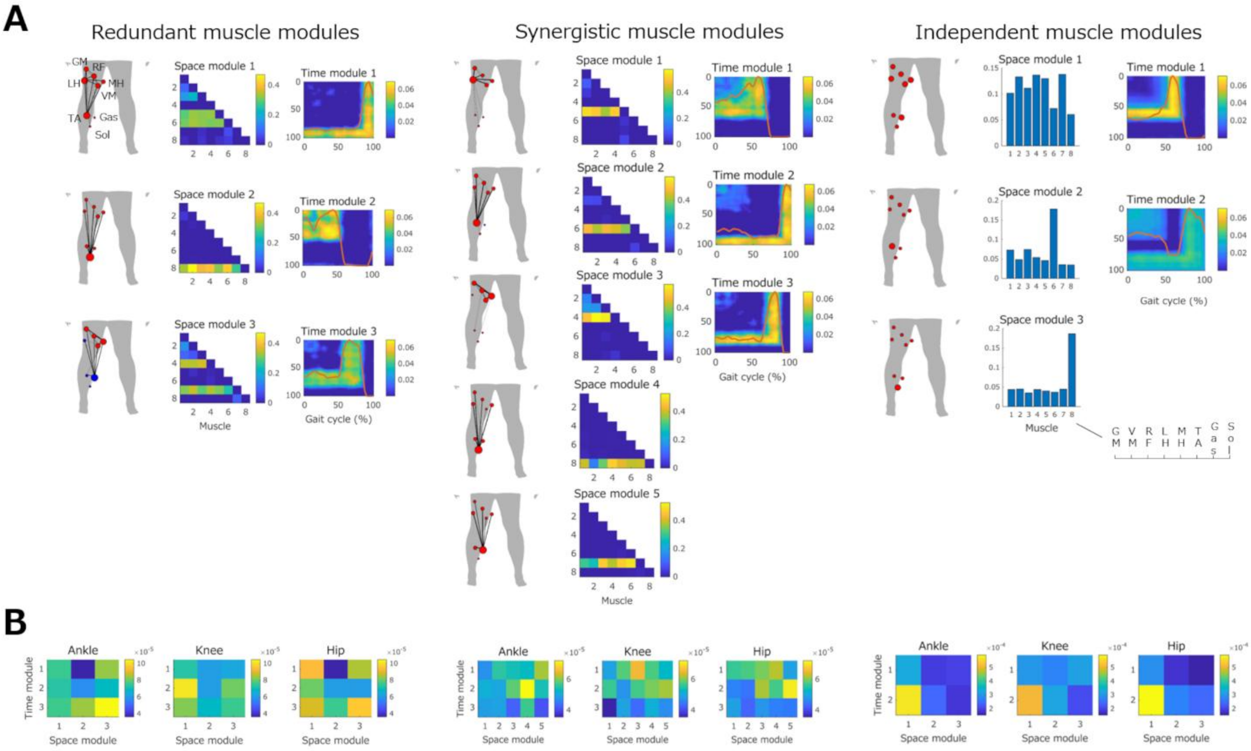


Supplementary Fig. 6. Functional muscle network modules in the paretic limb of participants with mild stroke

(A) Functional muscle network modules extracted from the paretic limb of participants with mild stroke. Node size indicates the relative strength of each muscle contribution, and line thickness indicates connectivity strength among muscles. Colored node groups denote subnetworks identified using link-based community detection. Space-module connectivity matrices show pairwise information strength among muscles. Time-module matrices show the temporal strength of information connectivity across the gait cycle. Muscle order was identical across all spatial modules: GM, VM, RF, LH, MH, TA, Gas, and Sol. (B) Relationships between space modules and time modules for each joint in the paretic limb of participants with mild stroke. Connectivity matrices show coupling strength between space and time modules for each joint. Individual space modules were associated with multiple time modules depending on joint-specific functional contributions. GM, gluteus medius; VM, vastus medialis; RF, rectus femoris; LH, lateral hamstrings; MH, medial hamstrings; TA, tibialis anterior; Gas, gastrocnemius; Sol, soleus.

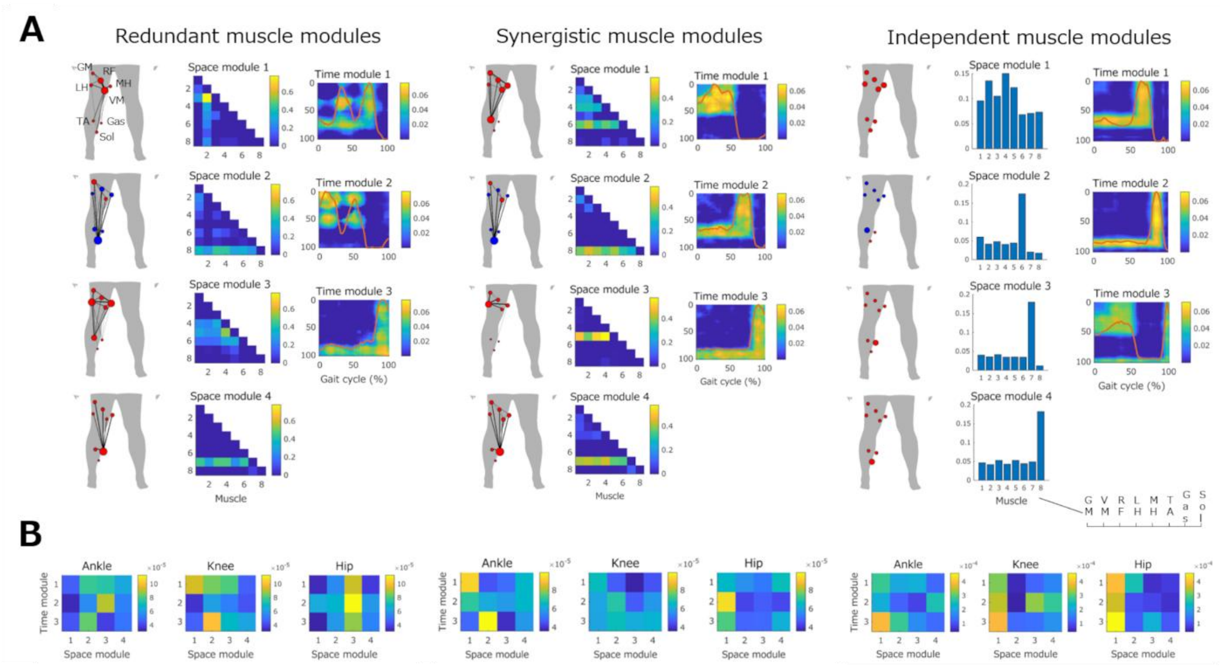


Supplementary Fig. 7. Functional muscle network modules in the paretic limb of participants with severe stroke

(A) Functional muscle network modules extracted from the paretic limb of participants with severe stroke. Node size indicates the relative strength of each muscle contribution, and line thickness indicates connectivity strength among muscles. Colored node groups denote subnetworks identified using link-based community detection. Space-module connectivity matrices show pairwise information strength among muscles. Time-module matrices show the temporal strength of information connectivity across the gait cycle. Muscle order was identical across all spatial modules: GM, VM, RF, LH, MH, TA, Gas, and Sol. (B) Relationships between space modules and time modules for each joint in the paretic limb of participants with severe stroke. Connectivity matrices show coupling strength between space and time modules for each joint. Individual space modules were associated with multiple time modules depending on joint-specific functional contributions. GM, gluteus medius; VM, vastus medialis; RF, rectus femoris; LH, lateral hamstrings; MH, medial hamstrings; TA, tibialis anterior; Gas, gastrocnemius; Sol, soleus.

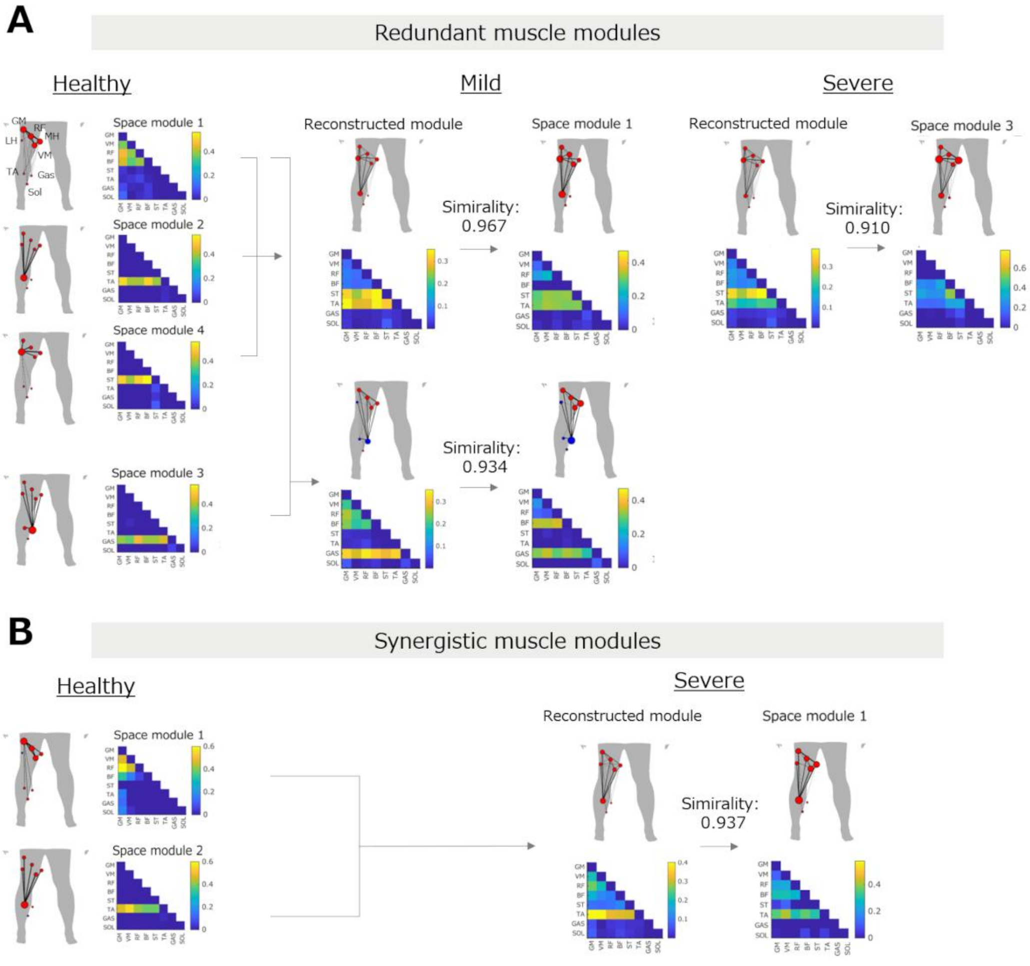


Supplementary Fig. 8. Severity-specific reconstruction of functional muscle network modules

(A) Reconstruction of redundant functional muscle network modules in the mild and severe stroke subgroups. Representative healthy spatial modules were merged using non-negative least squares to reconstruct the subgroup-specific spatial modules. Similarity values indicate the cosine similarity between the reconstructed and subgroup-specific spatial modules. (B) Reconstruction of the synergistic functional muscle network module in the severe stroke subgroup. Representative healthy spatial modules were merged using non-negative least squares to reconstruct the subgroup-specific spatial module. Similarity values indicate the cosine similarity between the reconstructed and subgroup-specific spatial module. GM, gluteus medius; VM, vastus medialis; RF, rectus femoris; LH, lateral hamstrings; MH, medial hamstrings; TA, tibialis anterior; Gas, gastrocnemius; Sol, soleus.

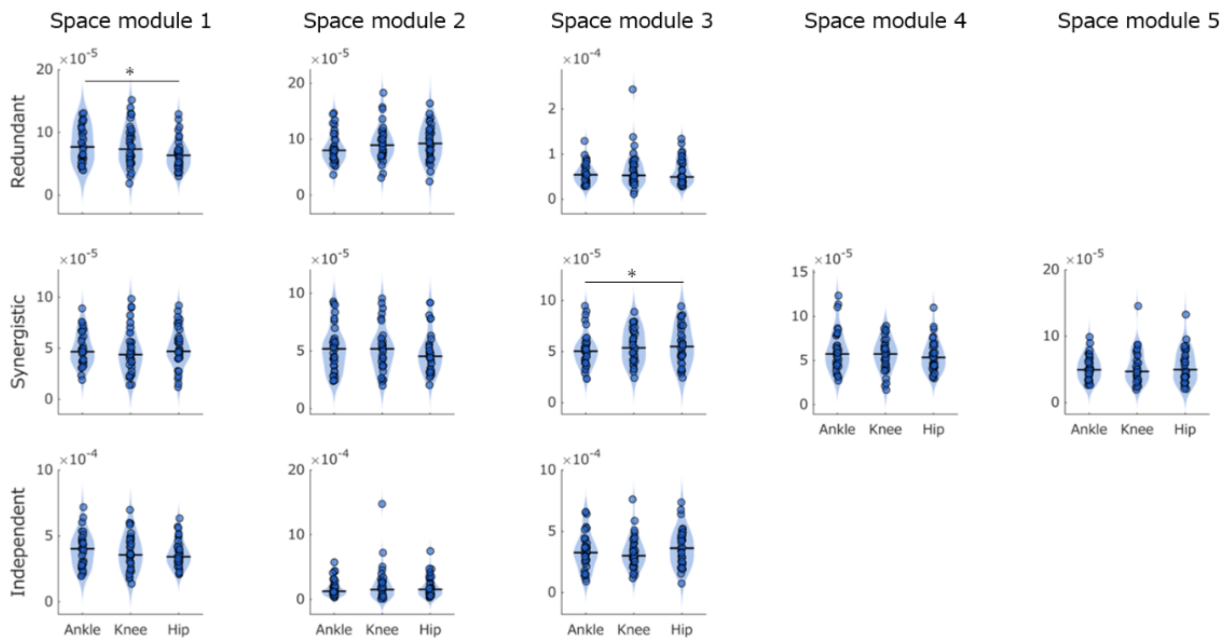


Supplementary Fig. 9. Joint-specific contributions of functional muscle network modules in the non-paretic limb of participants with stroke

Joint contributions were compared across redundant, synergistic, and independent space modules in the non-paretic limb of participants with stroke. Violin plots show the distribution of joint-specific contributions for the ankle, knee, and hip joints within each space module. Horizontal black lines denote median values. *Holm-adjusted $p < 0.05$

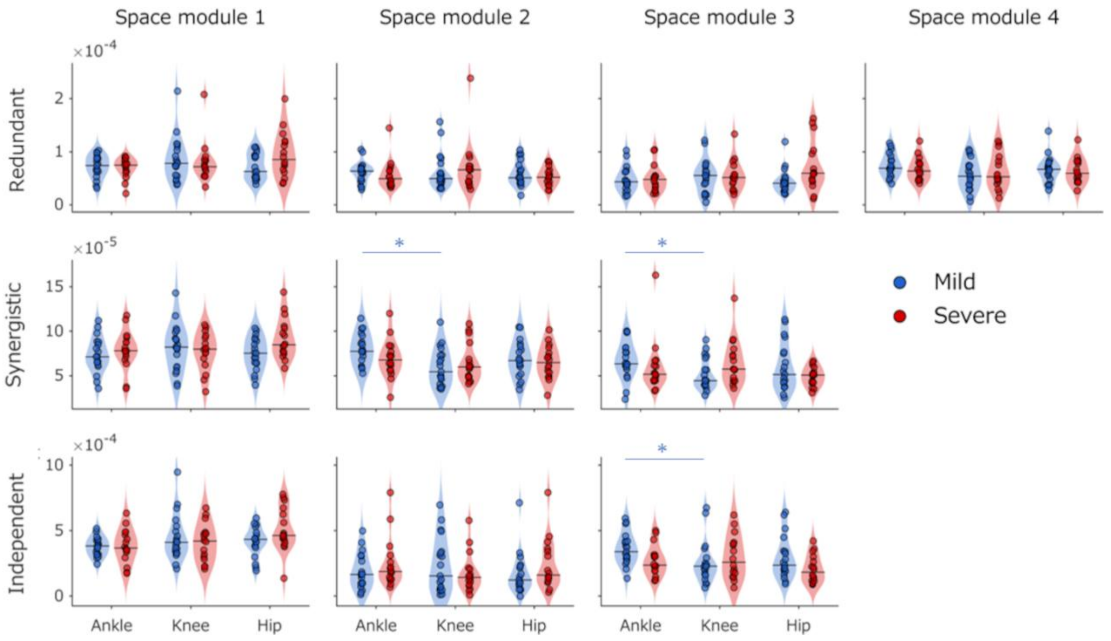


Supplementary Fig. 10. Severity-specific joint contributions of functional muscle network modules in participants with stroke

Joint contributions were compared across redundant, synergistic, and independent space modules in the mild and severe stroke subgroups. Blue and red violin plots denote the mild and severe subgroups, respectively. Violin plots show the distribution of joint-specific contributions for the ankle, knee, and hip joints within each space module, with horizontal black lines denoting median values. Blue brackets and asterisks mark significant post hoc differences between joints in the mild subgroup following significant Joint × Severity interactions identified by mixed-effects ART-ANOVA. * Holm-adjusted $p < 0.05$

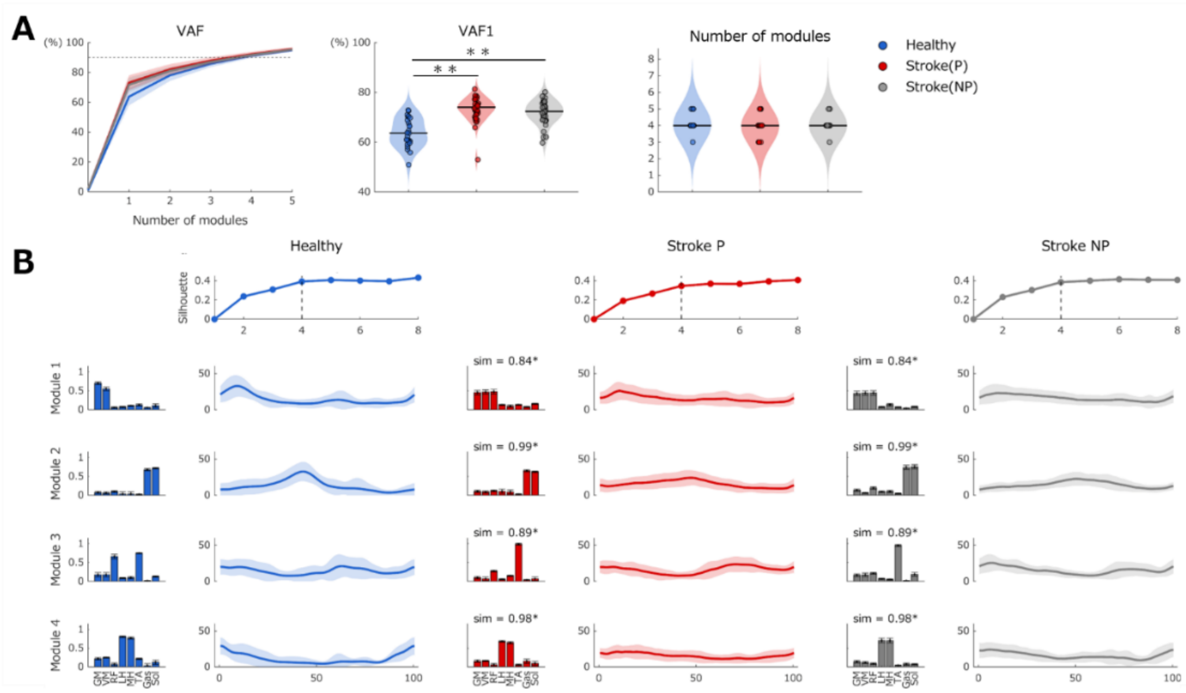


Supplementary Fig. 11. Muscle coordination patterns identified using NMF

(A) Group comparisons of total VAF, VAF1, and module number. (B) Silhouette scores and representative muscle synergy modules identified in each group. Bar graphs show spatial muscle weightings, and line plots show temporal activation patterns across the gait cycle. Waveforms are presented as mean ± 95% confidence intervals. Similarity between corresponding modules is reported as cosine similarity. Blue denotes healthy controls, red denotes the paretic limb of participants with stroke, and gray indicates the non-paretic limb. The similarity thresholds were 0.828 for the paretic limb and 0.830 for the non-paretic limb. **Holm-adjusted $p < 0.01$. GM, gluteus medius; VM, vastus medialis; RF, rectus femoris; LH, lateral hamstrings; MH, medial hamstrings; TA, tibialis anterior; Gas, gastrocnemius; Sol, soleus; NMF, non-negative matrix factorization.

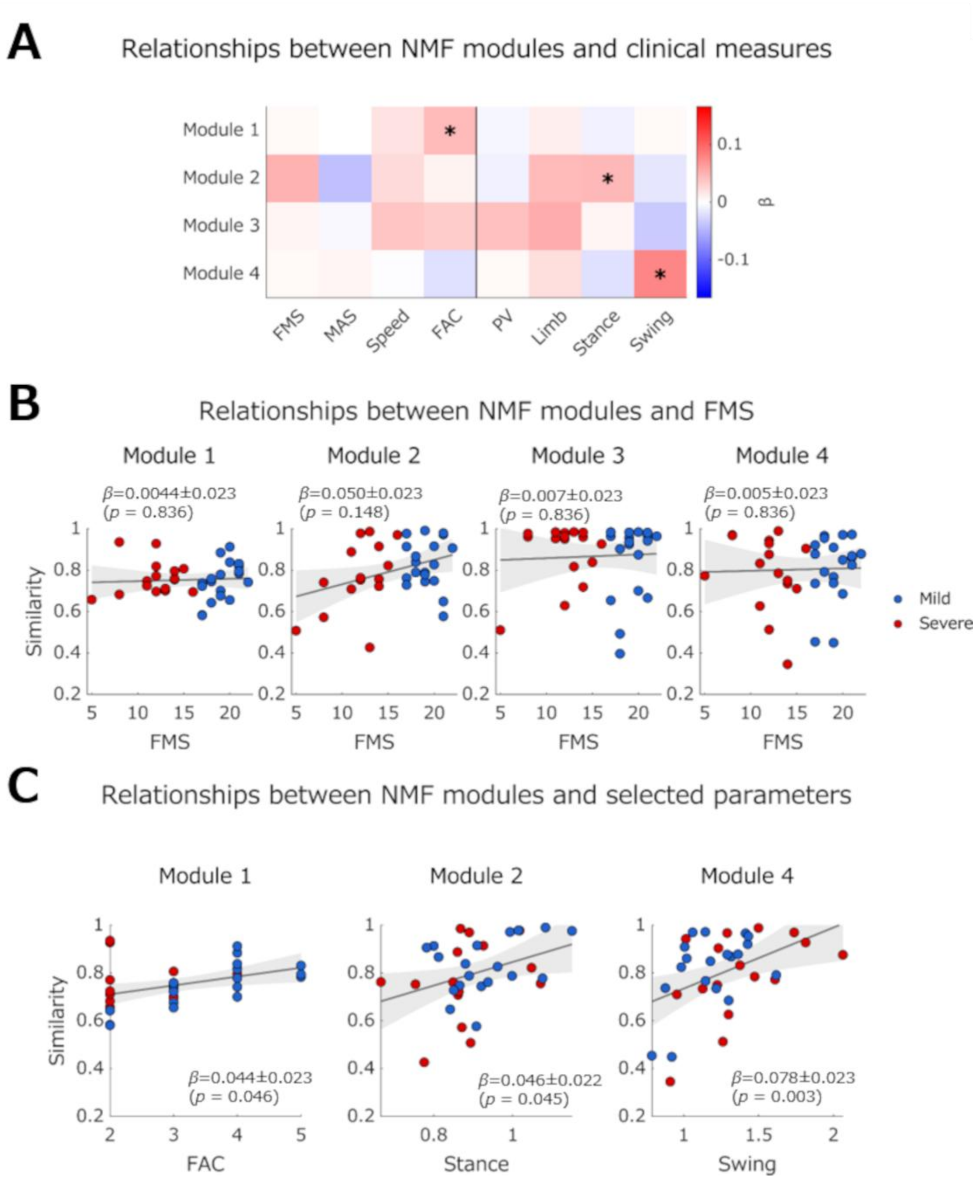


Supplementary Fig. 12. Relationships between NMF modules and clinical parameters

(A) Heatmap of regression coefficients (β) from mixed-effects models examining associations between NMF module similarity and clinical and gait-related measures. Associations with gait-related measures (PV, Limb, Stance, and Swing) were adjusted for FMS. (B) Associations between NMF module similarity and FMS. (C) Associations between NMF module similarity and selected gait-related measures. In panels B and C, blue indicates the mild subgroup and red indicates the severe subgroup. Solid lines and shaded areas represent the fitted regression lines and 95% confidence intervals, respectively. Regression coefficients (β) are presented as estimates ± standard errors. In panel A, *FDR-adjusted $p < 0.05$. FMS, Fugl–Meyer Assessment lower-extremity synergy scores; MAS, Modified Ashworth Scale; FAC, Functional Ambulation Category; PV, Planar Covariation; Limb, Limb angle range of motion; Stance, Stance time ratio; Swing, Swing time ratio; NMF, non-negative matrix factorization.

Supplementary Table 1. Severity-dependent associations between information components and clinical and gait-related measures

| | Redundant: Metric × Severity | | | | | Independent: Metric × Severity | | | | |
|---|---|---|---|---|---|---|---|---|---|---|
| | β_Mild ($\times 10^{-1}$) | β_Severe ($\times 10^{-1}$) | F | p | $\eta p^2$ | β_Mild ($\times 10^{-1}$) | β_Severe ($\times 10^{-1}$) | F | p | $\eta p^2$ |
| FMS | −0.054 | 0.077 | 0.933 | 0.683 | 0.029 | −0.030 | −0.073 | 0.120 | 0.731 | 0.004 |
| MAS | −0.040 | −0.004 | 0.193 | 0.663 | 0.006 | 0.056 | 0.021 | 0.232 | 0.633 | 0.007 |
| Speed | −0.020 | 0.108 | 3.094 | 0.176 | 0.090 | −0.001 | −0.066 | 0.863 | 0.360 | 0.027 |
| FAC | −0.028 | 0.131 | 4.931 | **0.041** | 0.137 | −0.005 | −0.099 | 2.084 | 0.158 | 0.063 |
| PV | −0.040 | −0.087 | 0.428 | 0.955 | 0.013 | 0.053 | 0.049 | 0.003 | 0.957 | 0.0001 |
| ROM | −0.022 | 0.055 | 1.272 | 0.267 | 0.039 | 0.011 | −0.068 | 1.594 | 0.270 | 0.048 |
| Stance | −0.029 | −0.043 | 0.047 | 0.829 | 0.001 | 0.025 | 0.005 | 0.118 | 0.830 | 0.004 |
| Swing | 0.036 | 0.022 | 0.039 | 0.844 | 0.001 | −0.026 | 0.008 | 0.272 | 0.845 | 0.010 |

Values are the estimated slopes (β) for the mild and severe subgroups, F statistics, $\eta p^2$, and Benjamini–Hochberg FDR-adjusted *p*-values for the Metric × Severity interactions. Bold values indicate statistically significant results. FDR, false discovery rate; FMS, Fugl–Meyer Assessment lower-extremity synergy scores; MAS, Modified Ashworth Scale; FAC, Functional Ambulation Category; PV, planar covariation; ROM, limb angle range of motion; Stance, stance time ratio; Swing, swing time ratio.

Supplementary Table 2. Post hoc comparisons of functional muscle network modules between the mild and severe subgroups

| | Module | Severe−Mild (×10$^{-5}$) | F | p | ηp2 |
|---|---|---|---|---|---|
| Redundant | 1 | 2.234 | 5.274 | **0.047** | 0.038 |
| | 2 | −0.588 | 0.366 | 1.000 | 0.002 |
| | 3 | 2.891 | 8.839 | **0.014** | 0.062 |
| | 4 | −0.212 | 0.047 | 1.000 | 0.0003 |
| Synergistic | 1 | 1.718 | 6.442 | **0.038** | 0.061 |
| | 2 | −0.292 | 0.185 | 0.667 | 0.002 |
| | 3 | −1.077 | 2.531 | 0.229 | 0.025 |
| Independent | 1 | 9.664 | 3.673 | 0.174 | 0.035 |
| | 2 | 7.326 | 2.119 | 0.192 | 0.021 |
| | 3 | −8.451 | 2.820 | 0.192 | 0.027 |

Values are the estimated difference in module strength between the severe and mild subgroups (Severe−Mild), F statistics, $\eta p^2$, and Holm-adjusted *p*-values. Bold values indicate statistically significant results.

Supplementary Table 3. Severity-dependent associations between functional muscle network modules and clinical and gait-related measures

| | Module | Metric | β_mild ($\times 10^{-5}$) | β_severe ($\times 10^{-5}$) | F | p | ηp2 |
|---|---|---|---|---|---|---|---|
| Redundant | 1 | FMS | −1.710 | −0.300 | 0.298 | 0.589 | 0.009 |
| | 1 | MAS | −1.217 | −1.422 | 0.011 | 0.917 | 0.004 |
| | 1 | Speed | 0.083 | −2.266 | 3.017 | **0.042** | 0.088 |
| | 1 | FAC | −0.273 | −1.598 | 0.870 | 0.584 | 0.027 |
| | 1 | PV | 0.896 | −0.078 | 0.498 | 0.485 | 0.015 |
| | 1 | ROM | 0.183 | −0.264 | 0.115 | 0.737 | 0.004 |
| | 1 | Stance | 0.205 | −2.196 | 4.553 | **0.041** | 0.128 |
| | 1 | Swing | −0.587 | 1.356 | 2.381 | 0.265 | 0.071 |
| Redundant | 3 | FMS | 1.411 | −0.158 | 0.332 | 0.589 | 0.010 |
| | 3 | MAS | 0.736 | −1.465 | 2.058 | 0.322 | 0.062 |
| | 3 | Speed | 0.429 | −2.366 | 3.881 | **0.042** | 0.111 |
| | 3 | FAC | 0.760 | −0.062 | 0.290 | 0.583 | 0.009 |
| | 3 | PV | 0.356 | −1.053 | 0.935 | 0.485 | 0.029 |
| | 3 | ROM | 0.334 | −0.616 | 0.473 | 0.737 | 0.015 |
| | 3 | Stance | 0.287 | 0.887 | 0.222 | 0.640 | 0.007 |
| | 3 | Swing | −0.612 | −0.834 | 0.029 | 0.865 | 0.001 |
| Synergistic | 1 | FMS | −0.007 | −0.626 | 0.155 | 0.696 | 0.005 |
| | 1 | MAS | 0.110 | −0.618 | 0.646 | 0.427 | 0.020 |
| | 1 | Speed | 0.620 | −0.548 | 1.963 | 0.182 | 0.056 |
| | 1 | FAC | 0.196 | −0.378 | 0.421 | 0.520 | 0.013 |
| | 1 | PV | 1.067 | 4.407 | 0.646 | 0.427 | 0.020 |
| | 1 | ROM | 0.622 | 0.021 | 0.575 | 0.453 | 0.018 |
| | 1 | Stance | −0.318 | −0.725 | 0.308 | 0.583 | 0.009 |
| | 1 | Swing | −0.046 | 2.767 | 0.160 | 0.692 | 0.005 |

Values are the estimated slopes (β) for the mild and severe subgroups, F statistics, $\eta p^2$, and Benjamini–Hochberg FDR-adjusted *p*-values for the Metric × Severity interactions. Bold values indicate statistically significant results.
